# A large-scale sORF screen identifies putative microproteins and provides insights into their interaction partners, localisation and function

**DOI:** 10.1101/2023.06.13.544808

**Authors:** Dörte Schlesinger, Christopher Dirks, Carmen Navarro Luzon, Lorenzo Lafranchi, Jürgen Eirich, Simon J Elsässer

## Abstract

The human genome contains thousands of potentially coding short open reading frames (sORFs). A growing set of microproteins translated from these sORFs are known to have important cellular functions. However, the majority remains uncharacterised. Thus, larger screens to find functional microproteins have become more vital. Here, we performed a high-throughput CRISPR/Cas9 knock-out screen with a customised library of 11,776 sORFs, curated from literature and databases to identify microproteins essential for cancer cell line growth. 16/17 tested candidates displayed a reproducible knockout phenotype. We selected our top six hits, consisting of 11 to 63 amino acids. Various of these candidates localised to distinct subcellular compartments and the majority showed specific interaction partners. Endogenous tagging demonstrated translation of an sORF in the CENPBD2P pseudogene that bears no resemblance to the CENPBD2P name-giving CENPB DNA binding domains. For two candidates, uORFs in the DSE and NUTF2 genes, the microprotein supplied *in trans* ameliorated the growth defect of the respective knock-out. RNA-seq analysis revealed however that gene expression changes in the knock-out could only partially be rescued. Overall, we identified various putative microproteins and a microprotein-producing pseudogene that might be involved in cancer cell growth, but also illustrate the limitations and caveats of sORF functional screening and characterisation.

## Introduction

Microproteins are polypeptides originating from short open reading frames (sORF) of less than a hundred codons. For a long time, they have been understudied as it is difficult to distinguish coding from non-coding sORFs. In recent years, the number of putatively translated sORFs could be narrowed down from hundred-thousands or millions to various thousands due to the advent of ribosome profiling and advances in bioinformatic and proteomic techniques. Consequently, efforts are now being made to include sORFs with robust translation evidence into databases such as GENCODE (1, 2). The field has been growing steadily, though it is still unclear how many functional coding sORFs there are and only relatively few microproteins have been characterised to this point. These arise from a variety of origins, including canonical protein-coding transcripts where microproteins can be translated for example as so-called upstream open reading frames (uORFs) from the 5’ “untranslated” region (3) or overlapping and out-of-frame with the canonical ORFs (4). Microproteins can also be produced from transcripts that were previously deemed non-coding such as long non-coding RNA (lncRNA) (5–8) and microRNAs (9). Additionally, sORF translation from pseudogenes has been demonstrated (10), though to our knowledge, no functional microprotein has been characterised from a pseudogene so far and indeed, pseudogenes are often excluded from large scale translation studies due to potential similarity with their canonical origin and difficulties in scoring them according to conservation criteria (11). Due to their small size, microproteins may be ideal for performing fine-tuning tasks (12). However, a number of characterised microproteins were also shown to carry out versatile or even essential cellular functions including roles in signalling (13–17), metabolism (18–24), and stress and repair pathways (25–28). Thus, to better understand cellular function, it is crucial to identify many more bioactive microproteins from the set of predicted coding sORFs. Yet, only few functional high-throughput screens of non-canonical ORFs have been performed to-date (29): three studies, screening several hundred sORFs, have been carried out in plants (30) or in yeast (31, 32) and more recently two screens have been performed in mammalian cells, surveying 2353 novel ORFs (33) and 553 non-canonical ORFs (34) respectively, while this study was ongoing. Here, we utilise a CRISPR/Cas9 drop-out screen with a human sORF-specific library of 11,776 sORF candidates to screen for microproteins required for growth and survival in three human cancer cell lines.

## Results

### An sORF-specific high-throughput CRISPR drop-out screen to identify sORFs required for cell survival

In order to design a comprehensive CRISPR library targeting human sORF candidates, we curated sORFs **(Fig. 1A)** from published Ribo-Seq, mass spectrometry (MS), bioinformatic or combined evidence studies (10, 11, 43–46, 35–42). We further included various publications describing individual microproteins (8, 22, 53, 25, 26, 47–52), UniProt reviewed human proteins below 60 amino acids in length and candidates from the sORF.org database (54, 55), applying the filtering criteria outlined in **Table S1** and the methods section. We used CRISPOR (56, 57) to design sgRNAs against the collected sORFs, requiring that each sORF had to be targeted by at least two sgRNAs **(Fig. S1A)**. For sORFs with more than 8 possible sgRNAs, we retained the 8 best guides. Our final screening set included 11,776 sORFs targeted by a total of 50,136 sgRNAs with a median of 6 sgRNAs/sORF **(Fig 1 B,C Fig. S1B)**. 2088 (17.73%) of targeted sORFs displayed overlap with canonical protein coding (PC) ORFs **(Fig 1D)**. Additionally, we included 292 positive control sgRNA targeting ribosomal genes, as well as 1000 non-targeting negative control sgRNA **(Fig. S2B)**.

**Figure 1:**
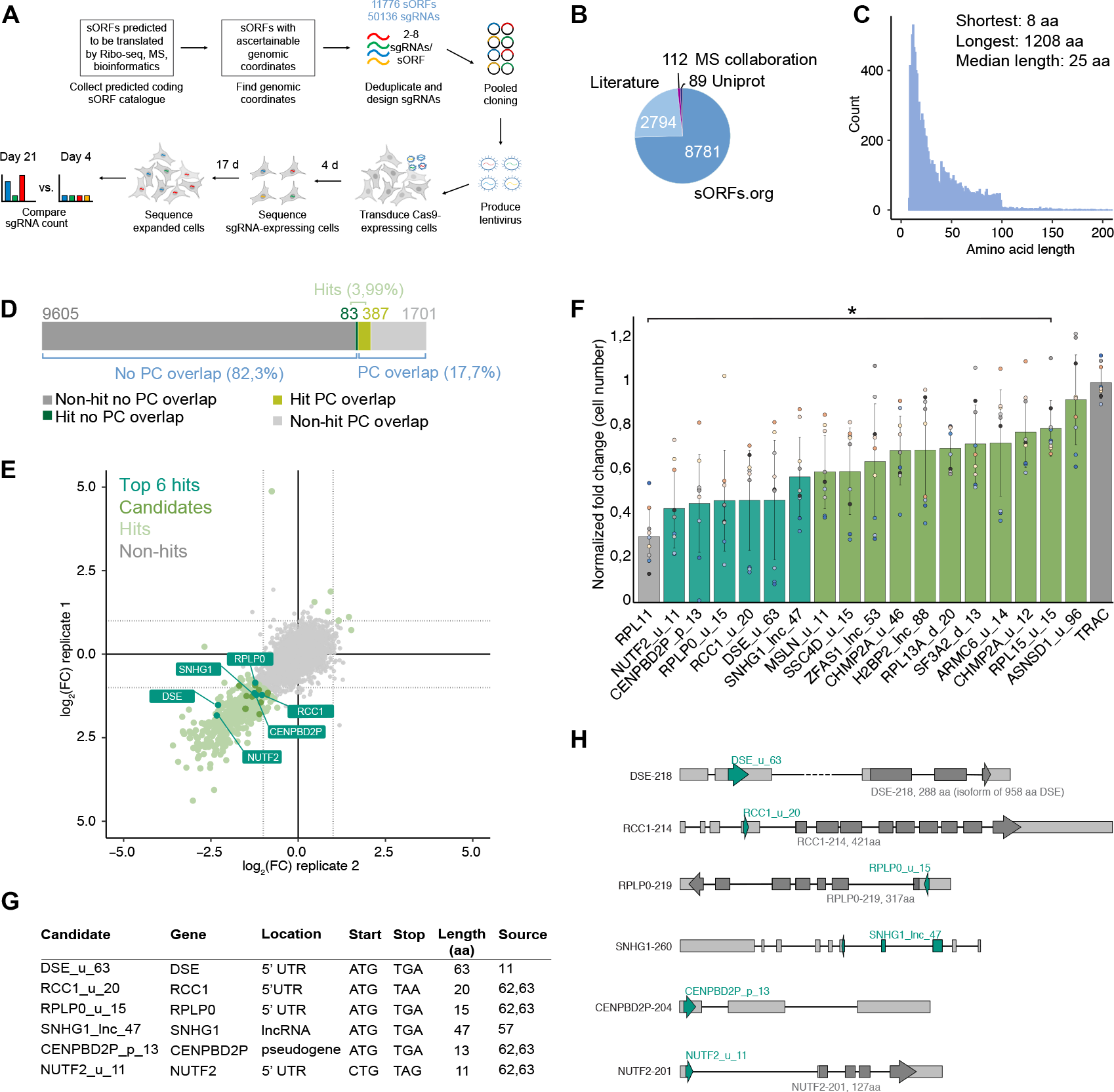
CRISPR screening to identify essential sORFs. A) Schematic of the sORF-specific CRISPR screen workflow. Cell schematics were created with BioRender. B) sORF catalogue by source of origin. C) Length diagram of ORFs in the CRISPR screen catalogue, the maximum 15 values were ommitted for visualization purposes. D) Stacked bargraph illustrating canonical protein-coding (PC) overlap of hit and non-hit sORFs, for PC overlap, any overlap (partial and full) was considered. E) Scatterplot summarizing the A375 10% FBS screen result, depicting log2 fold change (LFC) of the median of all sgRNAs in each replicate. Hits (light green): sORF with LFC (replicate average) ≥ 1 or ≤ –1. Candidates (green): filtered from hits based on genomic mapping, phenotypic evidence and lack of any overlap with protein-coding regions. Top 6 hits (dark green): selected from candidates based on results in F. F) Bargraph showing the results of the microscopy viability assay. A375 Cas9 cells were treated with the highest scoring screen sgRNA, a multi-guide positive (RPL11, grey) and a multi-guide negative (TRAC, dark grey) control. Fold change (FC) fixed cell number (Day 5 or Day 6)/live cell number (Day 1) was normalized to the average TRAC control FC of the respective experiment. n=9 with three independent experiments. Each data point is coloured according to replicate and experiment (yellow, blue, grey spectrum). Top 6 hits: dark-green, all other hits: green. Error bars represent standard deviation, * p-value < 0.05, compared to TRAC control and calculated by two-tailed unpaired student’s t-test, no asterisk indicates non-significance. G) Table summarizing the top six hits. H) likely Ensembl transcrips of origin for each of the top six candidates. dark green arrow: sORF, grey arrow: canonical ORF (if present), grey bars: exons.

The pool of 51,428 total sgRNAs was cloned and amplified into a lentiviral library, with which we carried out screens in three different cancer cell lines stably expressing Cas9 (A375 melanoma, HCT116 colon cancer, K562 leukaemia cells). We performed essentiality screens in either optimal serum (10% FBS in all cell lines), serum starvation conditions (1% or 0% serum in HCT116 and 1% serum in K562) or drug-resistant screening with 6-thioguanine (in A375, K562) **(Fig. S2A)**.

The A375 and HCT116 screens displayed a good dynamic range and strong correlation between replicates (correlation coefficient of 0.6-0.66 for sgRNAs and 0.75-0.8 for ORFs) **(Fig. S2A, B).** The A375 full serum screen showed the largest effect size and yielded a total of 470 hits with a log2 fold-change (LFC) smaller than -1 or larger than 1 (3.99%) **(Fig. 1D,E, Fig. S2E).** With few exceptions, the hit regions were located on medium to highly expressed transcripts, and most also showed Ribo-Seq signals **(Fig. S2F)**. Of note, 387/470 (82.34%) of these hits concerned sORFs that fully or partially overlapped with canonical protein coding (PC) ORFs, compared with only 17.73% PC overlap in the original library **(Fig. 1D)**. Hence, targeted regions were much more likely to show a growth phenotype when they were part of a canonical ORF. We considered that in these cases, it was difficult to discern loss of function effects of sORF versus ORF. Thus, we excluded PC overlapping hits. This yielded 83 remaining hits from which we manually curated 17 candidates for further validation by taking into account the supporting evidence of the sORF, strength and reproducibility of the phenotype, expression level, as well as the quality and number of sgRNAs **(Fig. 1E, Fig. S2 E)**. Of note, the fold-enrichment of these candidates was overall weaker than that of the positive controls and of many filtered sORF candidates with PC overlap. We reasoned that this could be due to sORFs displaying on average, much milder knockout phenotypes, compared to canonical proteins, which would be consistent with a potential fine-tuning function. All 17 candidates were also (mildly) downregulated in at least two replicates of any of the K562 viability screens and 15 out of 17 candidates displayed mild downregulation in the HCT116 screens, indicating that the majority of our candidates seems to display a similar phenotypic effect across the cell lines tested **(Fig. S2 D).**

We followed the same process with the HCT116 screens (full and serum starvation) but since this did not add any further candidates **(Fig. S2 C)**, we chose to perform follow-up experiments in A375 cells and systematically named our 17 candidates by their gene ID, location with respect to a canonical ORF, if present, or otherwise their transcript biotype, and putative amino acid length.

To validate the screening phenotype in an independent experiment, we turned to an imaging-based growth assay. To this end, we transfected A375-Cas9 cells with synthetic sgRNAs and followed cellular growth in a 96-well plate for 5-6 days. We chose the sgRNAs with the highest and most consistent effect size in the original screens to individually validate each of the 17 hits. As a positive control, we used a multi-sgRNA targeting the essential ribosomal gene RPL11, whereas as a negative control we utilised a multi-sgRNA targeting the non-expressed TRAC locus. For 16 out of 17 candidates, growth was significantly (p<0.05) reduced after transfecting the synthetic sgRNA, as compared to the TRAC control **(Fig. 1F).** We selected the six hits with the strongest phenotype for further studies **(Fig. 1 F, G, H).** Four of these top hits were uORFs (DSE_u_63, RCC1_u_20, RPLP0_u_15, NUTF2_u_11), one originated from a pseudogene (CENPBD2P_p_13) and one from a lncRNA (SNHG1_lnc_47) **(Fig. 1 G, H).** The candidate sORFs comprised between 11 and 63 codons and started with a canonical ATG start codon apart from NUTF2_u_11 **(Fig. 1G)**.

### CENPBD2P_p_13 is translated

Our CRISPR experiments suggested that the targeted sORF locations contained some functional element that elicited the reproducible growth defect. In a next step, we wanted to test whether our six predicted coding candidate sORFs were indeed translated. We did not observe any associated MS evidence of a sORF translation product in two tested published A375 datasets ((58) and Pride PXD016776) and one subcellular localization (59) dataset, though it also has to be noted that none of these data involved small protein enrichment. Further, we initially attempted to raise rabbit polyclonal antibodies, but limited possibility to select unique and highly immunogenic peptides in the short protein sequences did not allow us to derive selective and sensitive antibody sera. Hence, we resorted to an endogenous CRISPR knock-in strategy, fusing a GFP lacking a start codon to the C-terminal end of the respective endogenous sORF **(Fig. 2A)**. We expected that if the ribosome translates the sORF, it should also drive translation of the in-frame linker-GFP fusion. To carry out the knock-in, we used the microhomology-mediated end-joining based PITCh strategy, since it allows for very short homology arms (60). We co-transfected one plasmid containing the respective template and another containing the sORF-specific sgRNA, a linearizing sgRNA and Cas9 nuclease into A375 parental cells. For five out of the six chosen PITCh targets, transfection yielded GFP-expressing cells. We initially sorted the GFP-positive population and expanded the selectants. Despite the polyclonal nature of the resulting GFP-expressing lines, western blotting showed a single distinctive anti-GFP band for all candidates **(Fig. 2A)**. Targeting RCC1_u_20 resulted in an over 100 kDa large product, suggesting a preferential or exclusive off-target integration of the GFP cassette. However, four of the products fell into the expected molecular weight range of the desired sORF-GFP fusion. Encouraged by this, we sorted clones for each of the candidates to assess the respective integrations. The majority proved difficult to validate due to (off-target) insertions or complications in insert amplification **(Fig. S3A)**. Correct knock-in and functional translation could be confirmed only for CENPBD2P_p_13: Firstly, we performed a pulldown for GFP followed by mass spectrometry and were able to detect peptides tiling the microprotein-GFP-fusion in this assay **(Fig. 2B)**. Secondly, both parental and knock-in PCR products could be detected via genomic PCR indicating that the knock-in is heterozygous **(Fig. S3D).** Thirdly, the endogenous CENPBD2P_p_13-GFP knock-in clone showed a single western blot band at a similar height to an overexpressed CENPBD2P_p_13-GFP fusion control **(Fig. S3C).** Thus, we could confirm previous predictions that CENPBD2P_p_13 is indeed translated.

**Figure 2:**
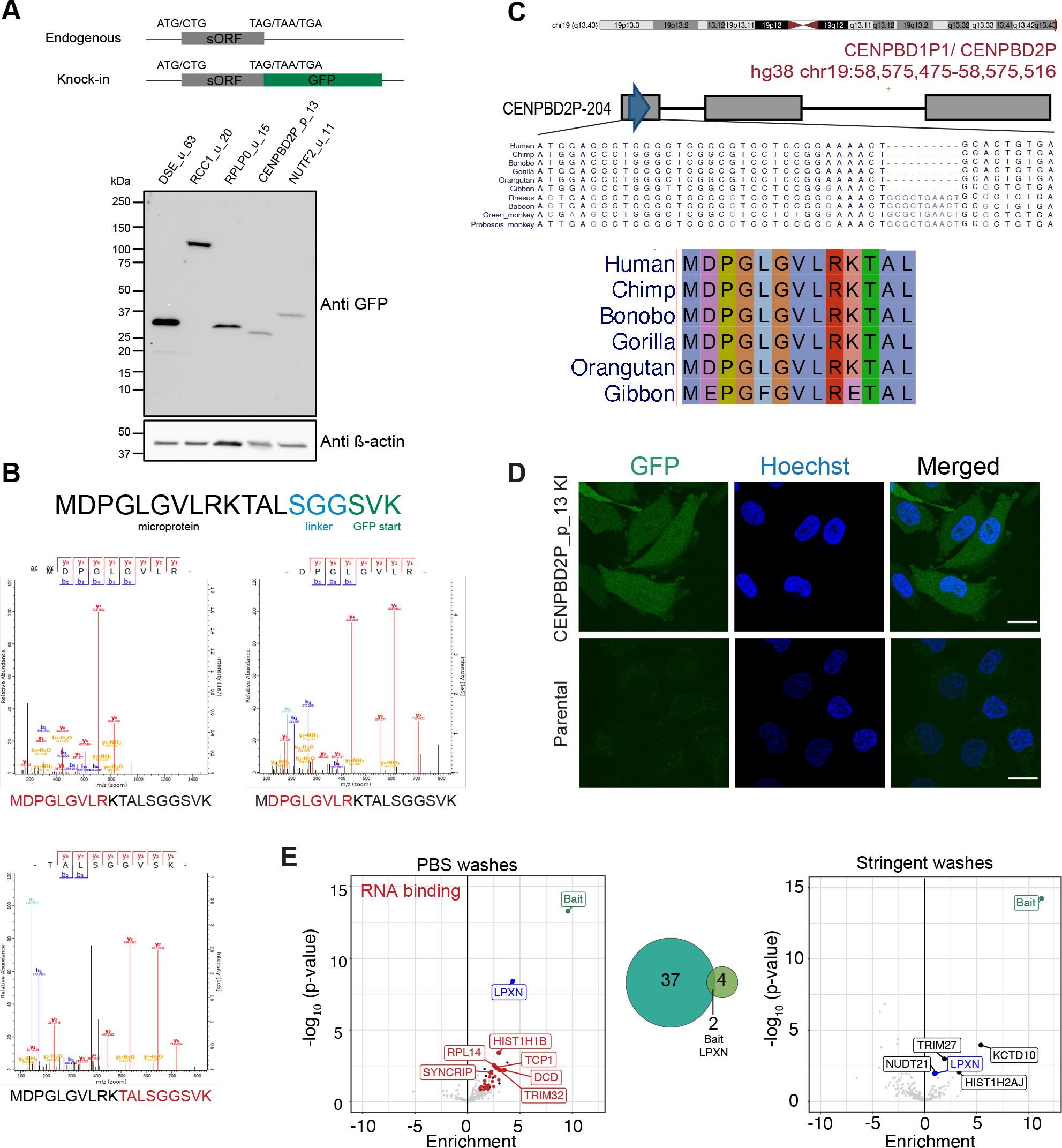
CENPBD2P_p_13 is translated. A) Schematic of knock-in strategy and Western blot of GFP+-sorted polyclonal knock-in A375 cells for each hit. The lower band in DSE_u_63 is likely a degradation product since it is smaller than full-length GFP. B) MaxQuant MS/MS spectra of the detected unique tryptic peptides for the CENPBD2P_p_13 fusion protein. Amino acid sequence of CENPBD2P_p_13, linker and the first three amino acids of GFP are indicated at the top. C) Chromosome location (red line), transcript location (blue arrow) and nucleotide and amino acid level Multiz 30 mammals alignment of CENPBD2P_p_13. Chromosome location, nucleotide alignment and phylogeny tree were outputted from http://genome.ucsc.edu. D) Live cell confocal microscopy of A375 monoclonal CENPBD2P_p_13-GFP knock-in cells and parental controls, with identifical laser and Fiji settings. green: CENPBD2P_p_13-GFP, blue: HOECHST 33342. Shown is a single plane from a z-stack. Scale bar represents 20 µm. E) Volcano plots and venn diagram showing results of mass-spec coupled GFP-trap CO-IP with either PBS washing conditions (right) or stringent washing conditions (left, see methods) of 2 biological and 2 technical replicates in each condition. Prey had to be present in 3/4 replicates and is coloured according to functional groups and overlap between conditions. Hit threshold: LFC ≥ 1, DEP adjusted p-value < 0.05.

### CENPBD2P_p_13 displays diffuse localisation and interacts with LPXN

CENPBD2P_p_13 is a primate-specific 13 amino acid small protein originating from the CENPBD2P pseudogene **(Fig. 2C)**. The endogenous CENPBD2P_p_13-GFP fusion showed diffuse subcellular localisation by confocal live cell microscopy **(Fig. 2 D).** To address a putative biological activity of the CENPBD2P_p_13 microprotein, we set out to identify possible interaction partners and performed MS-coupled co-immunoprecipitation (Co-IP). Parental A375 cells served as a control for the IP. A first pulldown with PBS washes yielded various RNA-binding proteins as well as the cell-adhesion and transcriptional regulator Leupaxin (LPXN) **(Fig. 2E)**. To increase stringency, we performed a second Co-IP with higher salt concentrations and detergent. This yielded LPXN as the only common interactor between both pulldowns **(Fig. 2E, Fig. S4B,C).** Given that CENPBD2P_p_13 is short and our fusion protein is not expressed at high levels (**Fig. S3B,C)**, these results are in agreement with the expectation that CENPBD2P_p_13 would have one or few direct interaction partners. However, Co-IP experiments reveal little about specificity and strength of interactions, hence additional experiments, such as reciprocal pull-downs could be useful to validate the interaction and determine through which regions CENPBD2P_p_13 and LPXN interact.

### sORF products display distinct localisation and bind to proteins involved in a variety of processes

To investigate the localisation and interaction partners of the five remaining candidates for which we were not successful in generating a knock-in, we created stable cell lines expressing a GFP-fusion of the respective microprotein using the PiggyBac transposase system (System Bioscience) in our A375-Cas9 screening cell line. Of note, we also attempted generating stable cell lines with HA-tagged constructs but were not able to detect any signal after antibiotic selection **(Fig. S5C)**. After confirming GFP expression of the transfected cell lines, we performed confocal microscopy. RCC1_u_20 and NUTF2_u_11 were diffusely distributed across the cytosol and nucleus **(Fig. 3 B,E)**, similarly to the linker-GFP itself **(Supp. Fig. 5A,B).** However, the remaining three candidates showed specific subcellular distributions: DSE_u_63 resembles mitochondrial localization, RPLP0_u_15 shows localization in line with the ER/Golgi and SNHG1_lnc_47 possesses a vesicular appearance **(Fig.3 A,C,D).** These localizations are especially notable since except a predicted transmembrane domain in SNHG1_lnc_47, none of the putative microproteins contain specific organelle motifs **(Supp. Fig. 4A, Supp. Fig. 7 A,B)**. We again performed MS-coupled Co-IP with the five stable cell lines and a A375-Cas9 control to find putative interaction partners of the GFP-fusion proteins. We calculated enrichments against the control and additionally filtered out proteins/apparent contaminants that were co-purifying with more than one GFP-fusion (these are shown in grey lettering **Fig. 3 B,D**; full graphs in **Supp. Fig. 6A)**. In accordance with their subcellular localization profiles, mitochondrial transport and ER proteins were co-enriched with DSE_u_63 and RPLP0_u_15 respectively **(Fig. 3A, C)**. The DSE_u_63 Co-IP also contained many components of the proteasome (32/62 proteasome complex and 15/20 proteasome core complex) **(Fig.3 A; Supp. Fig. 6B)**. Further studies will have to show whether this is due to the exogenously expressed DSE_u_63 being degraded by the proteasome or playing a functional proteasome related role. Other GO terms enriched in the candidate Co-IPs include transport and cholesterol binding (RPLP0_u_15) and cell adhesion molecule and guanyl nucleotide binding (NUTF2_u_11) **(Fig. 3C,E, Supp. Fig. 6B)**. RCC1_u_20 and SNHG1_lnc_47 displayed rather sparse interaction profiles **(Fig. 3B,D, Supp. Fig. 6B)**. In summary, Co-IP experiments indicated distinct protein-protein interactions of the selected sORF peptides.

**Figure 3:**
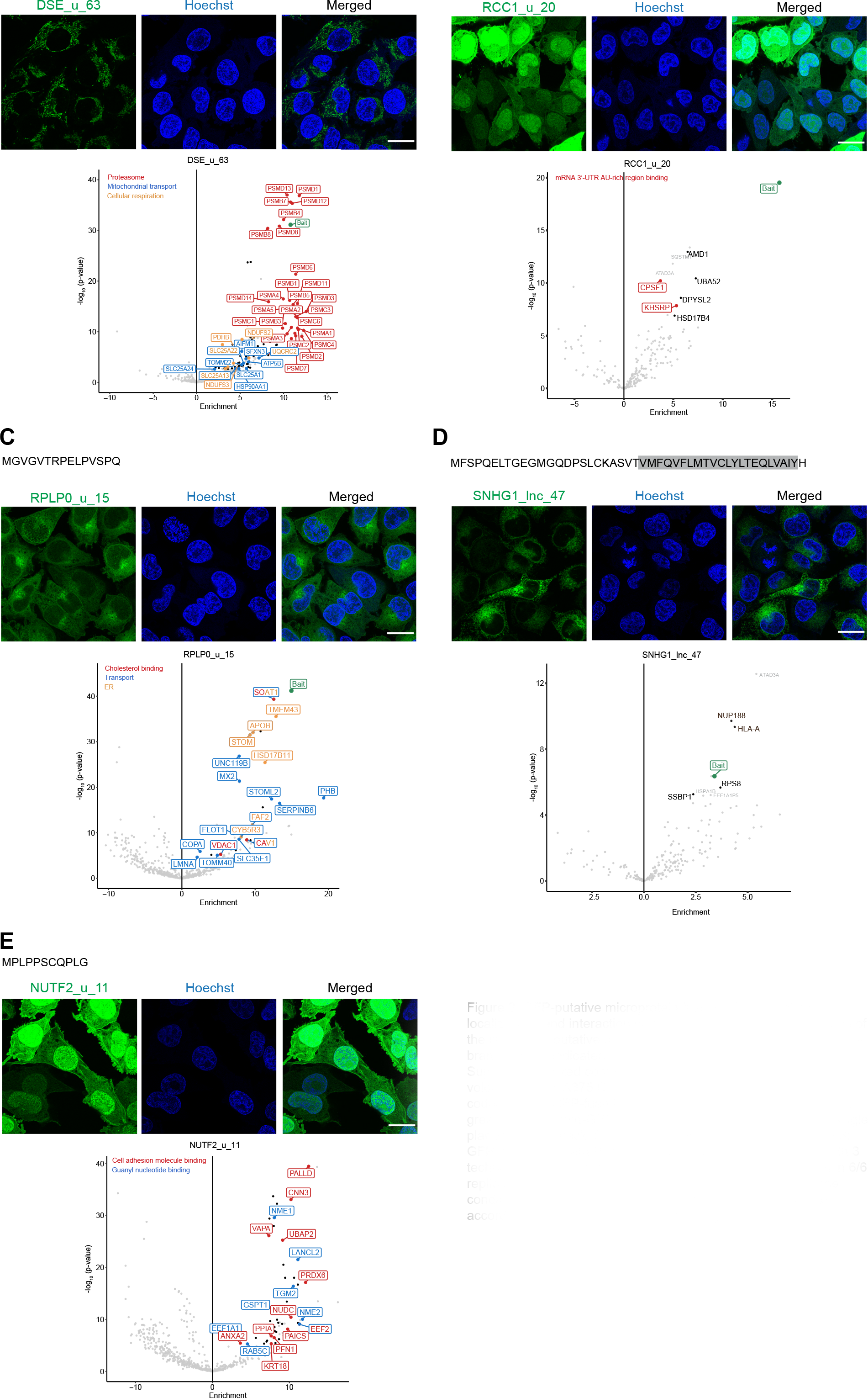
GFP-putative microprotein fusions display distinct localizations and interaction partners. (A-E) Amino acid sequence of the respective putative microprotein with ELM-predicted transmembrane domain indicated for SNHG1_lnc_47 (grey) in D (also see Sup Fig. 7B). Fixed confocal microscopy images and CO-IP MS volcano plots of A375 Cas9 cells stably expressing the respective codon-altered GFP-fused putative microproteins. For microscopy: green: microprotein-GFP, blue: HOECHST 33342. Shown is a single plane from a z-stack. Scale bar represents 20 µm. For Co-IP/MS: GFP-trap CO-IP with PBS washing conditions of 2 biological with 3

**Figure 4:**
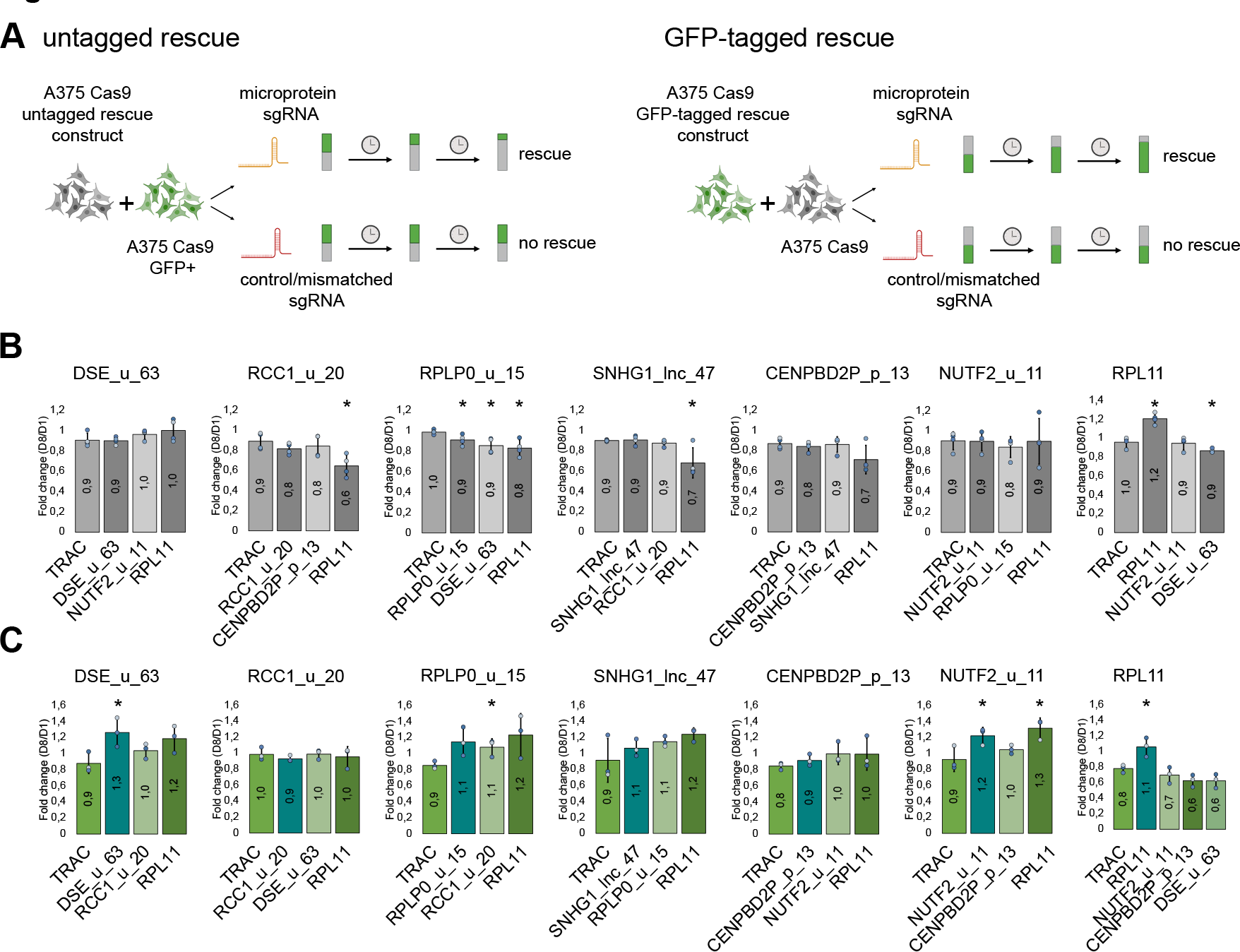
GFP-tagged but not untagged sORF product can mildy rescue the viability phenotype. A) Schematic of untagged (left panel) or tagged (right panel) co-culture rescue experiments. Cell schematics were created with BioRender. B) Barplots showing outcome of the untagged rescue experiment for each of the putative microprotein cell lines and RPL11 control cell line (right-most panel). A375 Cas9 cells stably expressing the respective untagged sgRNA-resistant microprotein candidate were co-cultured with parental A375 Cas9 cells stably expressing GFP. Each co-culture was treated with the negative control sgRNA (TRAC), the matched microprotein sgRNA and two mismatched sgRNAs (targeting a different sORF candidate and RPL11). Shown is the fold change (Day 8 /Day 1 after sgRNA-transfection) in the GFP-fraction measured by flow cytometry analysis. n=4 with 3 independent experiments, data points are coloured by experiment. C) Barplots showing outcome of the GFP-tagged rescue experiment for each of the putative microprotein cell lines and RPL11 control cell line (right-most panel). A375 Cas9 cells stably expressing the respective GFP-tagged sgRNA-resistant microprotein candidate were co-cultured with parental A375 Cas9 cells. Each co-culture was treated with the negative control sgRNA (TRAC), the matched microprotein sgRNA and two mismatched sgRNAs (targeting a different sORF candidate and RPL11). Shown is the fold change (Day 8 /Day 1 after sgRNA-transfection) in the GFP+ fraction measured by flow cytometry analysis. n=3 with 2 independent experiments (for SNHG1_lnc_47), n=3 with 3 independent experiments (for all others), data points are coloured by experiment. For both b) and c): All error bars indicate standard deviation. All p-values were calculated by two-tailed unpaired Student’s t-test compared to TRAC, *p<0.05, no asterisk indicates non-significance.

**Figure 5:**
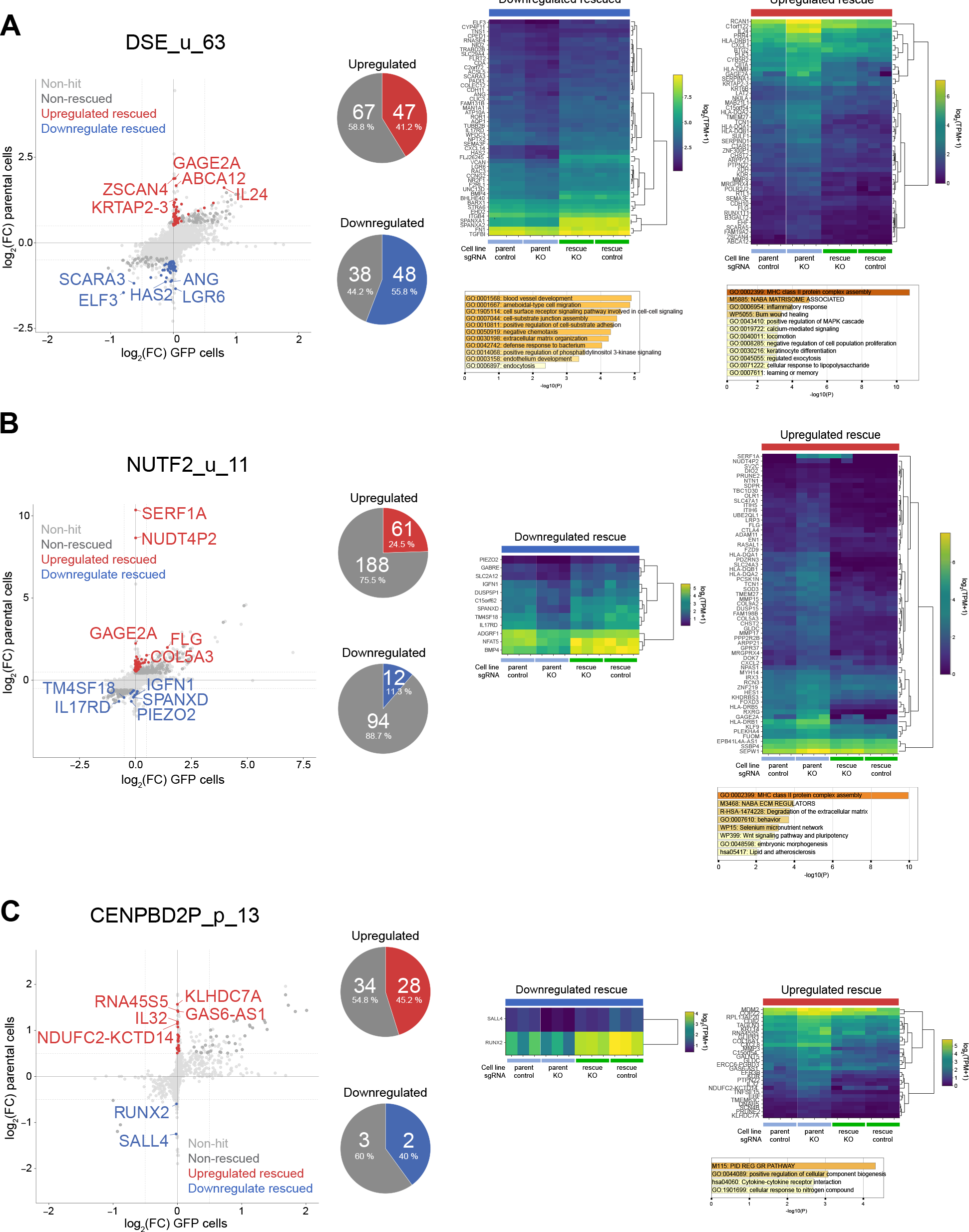
RNA-seq rescue experiments with GFP-tagged codon-altered putative microproteins. A-C) left panels: Scatterplot showing LFC expression levels (sORF-targeting sgRNA vs TRAC sgRNA) in GFP-tagged rescue A375 Cas9 cells and parental A375 Cas9 cells. Genes that were up-regulated or down-regulated in A375 Cas9 cells and rescued in microprotein-GFP expressing A375 Cas9 cells are indicated in red and blue respectively. Their fraction of the total altered genes is illustrated in a pie chart. middle panel: Heatmap of genes that were downregulated upon treatment of A375 Cas9 cells with the respective ORF sgRNA and that could be rescued in microprotein-GFP expressing A375 Cas9 cells. Bargraph of Metascape GO term analysis (where possible). right panel: Heatmap of genes that were upregulated upon treatment of A375 Cas9 cells with the respective ORF sgRNA and that could be rescued in microprotein-GFP expressing A375 Cas9 cells. Bargraph of Metascape GO term analysis. For all panels: Hits: genes displaying an average (of 2 replicates) LFC of ≥ 0.5 or ≤ –0.5 between microprotein sgRNA and TRAC sgRNA treated A375 Cas9 cells and showing an average expression variability of ≤10% between the TRAC conditions (microprotein-GFP vs. parental cells) and an DeSeq2 adjusted p-value of ≤ 0.05, Rescued hits: same as hits but additionally showing a difference in LFC of ≥ 0.5 between microprotein-GFP and parental cell lines.

### Complementation with GFP-tagged but not untagged microprotein partially rescues viability phenotype

Since specific subcellular localization and interactomes support a functional relevance of the putative sORF-encoded proteins, we sought to directly test if the microprotein expressed in *trans* could rescue a disruption of the endogenous sORF. This was especially important since five out of six sORF candidates were either located on transcripts implicated in cancer cell proliferation (61, 62) (RCC1_u_20, RPLP0_u_15, NUTF2_u_11, SNHG1), or possessed essential off-targets (RPS5 off-target for DSE_u_63 sgRNAs). We designed rescue constructs, in which codons were exchanged to their most common alternative. This made the exogenous sORFs resistant against sgRNAs targeting the endogenous locus and ensured that any rescue would be based on the encoded microprotein and not on a functional RNA sequence element. Placement of a selectable marker (Neo) under an IRES guaranteed expression of the sORF-encoding mRNA. After generating stable cell lines with the codon-altered constructs in a Cas9 background, we co-cultured the rescue cell lines with A375 Cas9 cells stably expressing GFP and analysed the single cell GFP+/GFP-ratio via flow cytometry (**Supp Fig. S8A)**. In case of rescue, cells expressing the exogenous microprotein (GFP-) should display a growth advantage over the control cells (GFP+) upon treatment with the matched sgRNA but not any of the mismatched sgRNAs **(Fig. 4A left panel)**. Indeed, a growth bias towards the rescue cells could be observed in the RPL11 positive control **(Fig. 4B, rightmost panel)**, however not for any of the top six sORF candidates **(Fig. 4 B)**. Since we had no means of detecting our untagged microproteins in this assay, the possibility remained that the sORFs were not efficiently translated from the supplied mRNA. Since microprotein-specific localization and interaction partners of our GFP fusion constructs suggested that the tag did not abrogate their function, we reasoned that expressing the respective microprotein-GFP fusions would allow us to interrogate a phenotypic rescue while being able to control for expression levels. We co-cultured GFP-tagged microprotein cell lines with A375 Cas9 parental cells, again assessing the single cell GFP+/GFP-ratio via flow cytometry **(Fig. 4 A right panel, Supp Fig. S8B)**. Here, a rescue effect should be apparent by an increase of the GFP+ population. Again, the positive RPL11 control construct was able to rescue its matched sgRNA condition **(Fig. 4C, rightmost panel)**. Additionally, we also observed a significant rescue of the respective sgRNA knockout phenotypes with GFP-tagged DSE_u_63 as well as NUTF2_u_11, but not any of the other microprotein candidates. Surprisingly both microproteins also showed a mild rescue (NUTF2_u_11) or rescue trend (DSE_u_63) upon treatment with RPL11 sgRNA. While we have not further dissected this, we speculate that this rescue could be due to the microproteins being involved in similar pathways as RPL11 or conferring a general proliferation/survival advantage that can help ameliorate the strong growth phenotype conferred by RPL11 targeting.

### Partial rescue of transcriptional perturbations in rescue cell lines

To further confirm the rescue, and to understand which cellular functions might be altered upon putative microprotein knockout, we wanted to employ a more sensitive readout and subsequently turned to RNA-sequencing experiments. We transfected both, the parental A375-Cas9 and microprotein-GFP fusion derivative cell lines, with the respective sORF-targeting or TRAC control sgRNA in triplicates and carried out RNA-seq analysis three days after transfection. We subsequently performed DESeq2 differential expression analysis contrasting target and TRAC sgRNA in each cell line background and identified differentially expressed genes (DESeq2 adjusted p-value ≤ 0.05, LFC ≥ 0.5 or ≤ -0.5) that were rescued in the respective microprotein-GFP expressing cell line (rescued by ≥0.5 LFC). We excluded highly variable transcripts (expression variability 10% or more amongst TRAC sgRNA controls).

We first turned our attention to the DSE_u_63 sORF: DSE_u_63 sgRNA treatment induced significant upregulation of 114 genes and downregulation of 86 genes **(Fig. 5A)**. Roughly half of these alterations (41.2% in the upregulated, 55.8% in the downregulated fraction) could be diminished in the DSE_u_63 GFP cell line, with rescued downregulated genes being enriched in cell migration and adhesion GO terms. Cell adhesion and signalling was also enriched in the rescued upregulated fraction, although the top scoring terms were associated with antigen processing and presentation. Hence, the deregulated gene sets did not provide an obvious mechanistic link to the apparent mitochondrial localization of the DSE_u_63 GFP fusion microprotein **(Fig. 3A)**. Nonetheless, our RNA-seq analysis corroborated that the DSE_u_63-GFP microprotein fusion supplied in *trans* could phenotypically and transcriptionally rescue an endogenous sORF knockout. We also inspected the gene sets not rescued by the microprotein-GFP fusion: the leading upregulated GO terms were p53 downstream effects **(Fig. S9A)**. We attributed this to sgRNA-induced genotoxic effects independent of sORF-targeting and/or off-target effects. Indeed, one of the DSE_u_63 sgRNA’s predicted off-targets is the essential gene RPS5 and upon inspection, we did indeed see downregulation of the RPS5 transcript (**Fig. S10C,D;** not visible in **Fig. S9A** since it was filtered due to variability between TRAC control conditions).

Next, we investigated the effects of NUTF2_u_11 knockout and whether they could be ameliorated in the respective GFP fusion cell line **(Fig. 5B)**. Treatment with the NUTF2_u_11 sgRNA induced upregulation of 249 and downregulation of 106 genes. 61 and 12 of these respectively were rescued by the NUTF2_u_11-GFP fusion. GO analysis did not yield any enrichment for the rescued downregulated fraction. The rescued upregulated fraction was enriched in ECM-connected GO terms, which could fit with the cell adhesion associated NUTF2_u_11 interactors (Fig. 3D). Additionally, MHC class II complex assembly associated genes were enriched, similarly to DSE_u_63 **(Fig. 5B)**. Indeed, some overlap existed in the rescuable upregulated genes between DSE_u_63 and NUTF2_u_11, including several in the HLA-D locus (**Fig. S10 A,B)**. While this explained the common MHC GO term, the source of this shared set of differential genes remained unclear. We considered that the TRAC sgRNA used as control for all experiments could target and downregulate these loci, but they were not amongst obvious TRAC off-targets predicted by CRISPOR (56, 57). Conversely, a large number of up- and downregulation events in the NUTF2_u_11 knockout could not be rescued, and these were enriched in GO terms relating to ribosome biogenesis, various signal and stress pathways (**Fig. S9B)**. Since NUTF2_u_11 is located upstream of a canonical ORF encoding the essential nuclear transport protein NUTF2, we wondered if our sORF-targeting sgRNA also perturbed the canonical NUTF2. Indeed, the NUTF2 RNA was one of the downregulated transcripts, which could not be rescued by the NUTF2_u_11-GFP microprotein fusion (**Fig. S9B)**. This suggested that targeting the NUTF2_u_11 uORF caused destabilisation of the entire transcript. A reduction of NUTF2 expression may in turn affect nuclear transport and indirectly processes that depend on it, such as ribosome biogenesis. This highlights the difficulty separating uORF function from that of the transcript or the canonical ORF product, especially if the canonical protein is also essential.

Finally, we also tested a rescue of CENPBD2P_p_13 knockout, given that we observed endogenous production of the CENPBD2P_p_13 microprotein **(Fig. 2)**. 62 genes were significantly upregulated, and 5 genes were significantly downregulated upon CENPBD2P_p_13 sgRNA-mediated knockout. Of these, 28 and 2 genes could be restored respectively in the CENPBD2P_p_13-GFP rescue cell lines **(Fig. 5C)**. GO terms did not reach high significance, given the small number of differential genes rescued. Upregulated non-rescued genes were enriched in p53 downstream pathways as in the case of DSE_u_63 above **(Fig. S9C).** Interestingly, the CENPBD2P pseudogene itself was among the non-rescued down-regulated genes (**Fig. S9C)**, again suggesting that targeting the sORF can affect transcript stability. In the case of canonical ORFs, nonsense-mediated decay is thought to be responsible for a downregulation of the transcript. In the light of this and the fact that CENPBD2P_p_13 microprotein in *trans* cannot rescue the growth defect, we speculate that the translation of CENPBD2P_p_13 sORF may be required to stabilise the CENPBD2P transcript. Interestingly, Chen et al. (2020) targeted a different CENPBD2P sORF downstream of CENPBD2P_p_13 and also observed a mild growth defect (33), further corroborating that the CENPBD2P pseudogene transcript may possess a non-coding function in cell growth.

Overall, RNA-seq analysis revealed sORF-knockout specific gene perturbations, and expression of the microprotein in *trans* rescued these gene expression changes to a variable extent. Gene sets perturbed downstream of the putative microproteins did not yield an immediate mechanistic hypothesis for how the putative microprotein may work, and also did not have a clear relation with the microprotein localization and protein interactions determined **(Figure 3)**.

In summary, though diverse assays used to pin down a possible cellular function did not converge on a coherent mechanistic picture, studying multiple microprotein candidates from a large-scale CRISPR screen exclusively targeting sORFs, we find unique cellular localizations, interactomes and transcriptional perturbations associated with the putative microproteins.

## Discussion

In the human genome, sORFs with coding potential exceed canonical protein-coding ORFs by 1-2 orders of magnitude. Ample evidence exists for their pervasive translation but the extent to which sORF-encoded microproteins contribute to cellular function is still unknown. Here, we carried out a sORF-specific large scale functional CRISPR/Cas9 screen targeting 11,776 sORFs with predicted coding potential from which we selected candidates to study for a more detailed characterization. Two other CRISPR screens targeting non-canonical ORFs in human cells have recently been published (33, 34). Prensner et al. 2021 also surveyed A375 cells, amongst others, and identified 15 non-canonical ORFs, knockout of which led to a growth phenotype, with a similar effect size (log2-fold change ≤ -1) to our screen (34). The proportion of hits was considerably higher compared to our study, given that their library targeted only 553 ORFs. The ORFs included in the latter study were larger on average (median length 74 aa, minimum 23 aa). While 50 out of these 553 sORFs were also contained in our library, only one relatively long sORF, ASNSD1_u_96, was a common hit between our studies. In our hands however, the growth phenotype of ASNSD1_u_96 was not reproduced in A375 cells in a screen-independent growth assay (**Fig. 1F**). The ASNSD1_u_96 microprotein has since been described as a component of the PAQosome (63). None of our other 16 candidates was included in the 553 ORFs (34). Conversely, our library contained only one of the other 14 downregulated ORFs in the Prensner screen ((34): TCONS_I2_00007040, this screen: PL01912), targeting of which did not score any phenotype in our study. Another CRISPR screen targeting 2353 ORFs was also reported, albeit not in A375 cells (33). The hits further characterised from this screen did not overlap with our hits.

The annotation of sORF has tremendously improved in the past years: a large community effort collected a set of 7264 sORFs with high-confidence Ribo-Seq signals for submission to GENCODE (2). SNHG1_lnc_47 is the only top hit included in this catalogue (now uniprot predicted protein A0A024R548). CENPBD2P_p_13 and RPLP0_u_15 were smaller than the 16 amino acid consortium threshold, but were present respectively in two and three of the published Ribo-seq datasets utilised by the consortium.

RCC1_u_20 and RPLP0_15 were originally extracted from the Ribo-seq-based sORF database sORFs.org (54, 55), with the latter currently showing some level of evidence in 30 different repository datasets. While RCC1_u_20 did not display any notable subcellular localisation nor interaction partners, RPLP0_u_15 expressed as a GFP fusion resembled Golgi/ER localisation, despite its short 15-amino acid sequence. Another hit with a distinct localisation is the putative 47-amino acid microprotein SNHG1_lnc_47, extracted here originally from a Ribo-seq-based study (35). SNHG1_lnc_47 seemed to localise to vesicles when fused to GFP and contains a predicted transmembrane domain. Functional follow up for either of these putative microproteins proved difficult since neither an untagged protein nor the GFP fusion could rescue the growth phenotype elicited by the respective microprotein sgRNA and we were not able to independently verify microprotein translation.

In contrast, through a GFP-knockin, we were able to validate CENPBD2P_p_13 translation, which initially was derived from the sORFs.org database (54, 55). CENPBD2P_p_13 originates from the CENPBD2P pseudogene previously categorized as non-coding and showed diffuse subcellular localization and scarce protein-protein interactions. The inability of the CENPBD2P_p_13 microprotein to rescue a sgRNA-mediated knockout in *trans* and the lack of CRISPOR-predicted viability-related sgRNA off-targets suggested that the growth defect observed in the original screen may relate to a non-coding function of the CENPBD2P pseudogene itself. In support of this, when assessing transcriptional consequences of CENPBD2P_p_13 sgRNA treatment, we observed downregulation of the CENPBD2P transcript, which could not be rescued by the microprotein expressed in *trans*. (33) We speculate that CENPBD2P_p_13 evolved as a regulatory sORF, with its translation boosting levels of the underlying transcript.

The DSE_u_63 sORF originates from a bioinformatics study mining genomes for evolutionary signatures of protein coding sORFs (11), but in contrast to most of our sORF library, no prior Ribo-Seq or mass spectrometry evidence exists for the DSE_u_63 microprotein. While the sORF showed one of the strongest growth defects initially, interpretation of the phenotypic and RNA-seq experiments were complicated by a CRISPR off-target effect towards a ribosomal protein, Rps5. All possible DSE_u_63 sgRNAs share the same off-target region within Rps5 and this is related to the evolutionary origin of this sORF. DSE_u_63 appears to have arisen in the primate lineage after the branching of macaques and gibbons, by a de-novo integration of a new chunk of DNA with high homology to part of the Rps5 coding sequence, reversed with respect to its original orientation in the Rps5 gene. The full DSE_u_63 sORF including start and stop codon can already be found in the macaque Rps5 mRNA gene, running antisense to the Rps5 coding sequence. Hence, the DSE_u_63 sORF appeared immediately through the inverse integration of the Rps5 fragment into the DSE gene. It has acquired only five sense mutations up to the hominid lineage. Thus, a mitochondrial localization and interactome of DSE_u_63 is unlikely to have evolved since its birth in the primate lineage - instead we hypothesise that by pure serendipity, a sequence encoded antisense to the Rps5 coding region was capable of producing a mitochondrially localised microprotein. Since the human microprotein has acquired five mutations as compared to the gibbon homolog, it would be interesting to experimentally test if the original sequence showed the same subcellular localization and putative phenotype, or both evolved further in the higher hominid lineage.

Finally, the shortest of our candidates is the putative 11 amino-acid microprotein NUTF2_u_11, also extracted from sORF.org (54, 55) **(Fig. 1G)**. While in the majority of datasets in this repository, this sORF is described as encoding a 11 amino acid microprotein, an alternative 13 codon version is also possible **(Fig. S2G)**. NUTF2_u_11 did not show a specific subcellular localization but displayed interaction partners enriched in cell adhesion terms. The growth defect of the NUTF2_u_11 sgRNA could be significantly rescued with a NUTF2_u_11 microprotein-GFP fusion expressed in *trans,* albeit only a minority of transcriptomic changes could be reverted. As with CENPBD2P_p_13, we observed downregulation of the underlying transcript as a result of targeting the NUTF2_u_11 sORF. This underscores the difficulty of separating sORF activity from other functional elements of the same transcript and the need for suitable rescue experiments.

For studying our microprotein candidates, we turned to a GFP fusion. We resorted to this approach because microprotein fusions with smaller tags, specifically HA and FLAG, which we generated in parallel, could not be detected. This suggests that the native microproteins may be suboptimally expressed from our plasmids, potentially lacking regulatory sequences that boost translation of the endogenous sORF. It is also likely that a well-folded GFP can protect a microprotein from degradation and hence boosts its steady state levels. This may explain why we only observed functional rescue in *trans* when we expressed the transgene microprotein as a GFP fusion. At the same time, we are aware that adding a large tag may disrupt native microprotein-protein interactions and hence the lack of strong interaction partners for some of the microproteins may reflect a perturbation of localization or interaction partners by GFP. Interesting to us was the fact that several small microproteins could localise to specific compartments even in the context of a GFP tag. Thus, in the absence of specific antibodies, finding the right tag for microprotein studies remains a major challenge in the field.

In summary, we identified six candidate sORFs and tested interaction partners and localizations for their encoded putative microproteins. Rescue experiments implicated two of these putative small proteins in cell proliferation, and RNA-seq experiments demonstrated a partial rescue of gene expression perturbations for three candidates, though for all tested hits, sORF knockout also induced knockdown of an off-target or the transcript of origin with functions of its own. We could demonstrate endogenous translation of the CENPBD2P_p_13 microprotein and our data suggests that its pseudogene of origin, CENPBD2P, is involved in cell proliferation. Our work contributes to the growing number of putative microproteins by adding characterization of various sORF candidates and translation evidence of a pseudogene-derived microprotein.

### Data Availability

Mass spectrometry primary data has been deposited on MassIVE under doi:10.25345/C5DZ03B70. RNA-seq primary and processed data has been deposited on GEO under GSE232375.

## Acknowledgements

CRISPR screening was carried out at the SciLifeLab CRISPR Functional Genomics unit (CFG) at Karolinska Institutet, funded by Science for Life Laboratory. CRISPR and genomics analyses were processed on the Uppsala Multidisciplinary Center for Advanced Computational Science (UPPMAX) provided by the National Academic Infrastructure for Supercomputing in Sweden (NAISS) under projects NAISS 2023/6-19, NAISS 2023/22-84, SNIC 2022/6-14. NAISS is funded by the Swedish Research Council through grant agreement no. 2022-06725. We thank Dr. Rui Branca for sharing a list of unannotated peptide candidates from proteogenomics experiments. We thank Prof. Iris Finkemeier and Paulina Heinkow for mass spectrometry service. We thank the BIC facility for enabling usage of their AiryScan microscopy, the Fernandez Capetillo group for allowing us to use their In Cell microscope, the Bartek and Lemmens groups for access to the Nikon Eclipse Ti2.

## Materials and Methods

### sORF catalogue assembly

Microprotein candidates were curated from literature (8, 10, 37–46, 11, 47–53, 64–66, 13, 67, 68, 14, 22, 25, 26, 35, 36) collaborative mass spectrometry data (69, 70), Uniprot (71) and the sORF.org database (54, 55) with the criteria described in **(Table S1)**. Each sORF was assigned a number with the first two letters indicating the peptide source (PL: literature, PM: mass spectrometry, PS: sORFs.org database, PU: Uniprot). The resulting sORF catalogue was deduplicated by their amino acid sequence. Since genomic coordinates were absent in some publications and differently reported between studies, we mapped all candidates to the human genome (GRCh38) using Scipio (72) based on the amino acid sequence supplied in the original source. Since we noticed that searching for a start codon led to errors (partially due to some sORFs containing non-canonical start codons), we first removed the starting Methionine, carried out the mapping and then expanded the mapped interval again by three nucleotides on the 5’ end. In cases where both amino acid sequence and genomic coordinates were provided, conflicts were resolved by keeping Scipio results when at least 80% of the Scipio prediction overlapped with the original annotation, and retaining genomic coordinates supplied in the original source otherwise. The mappable sORFs were then deduplicated once more and subsequently carried through to sgRNA design.

### CRISPR screening

#### sgRNA design

To design sgRNAs against each of the sORFs in the catalogue, we utilised the CRISPOR design tool (56, 57) specifying the sORF coordinates with an additional 20-nucleotide flanking region. We excluded sgRNAs within the lowest 20% of all cutting frequency determination (CFD) scores, as well as sgRNAs displaying more than four thymidines in a row **(Supp. Fig. 1A)**. If the sORF possessed only one targeting sgRNA, the sgRNA and sORF were dismissed from the library, while all sgRNAs were kept if the sORF could be targeted by two to eight sgRNAs. The CFD specificity score and the Doench et al. efficiency score (73) supplied by CRISPOR were each normalised to a 0 to 1 value range each (x - min(scores)) / (max(scores) - min(scores)) and subsequently added to create an aggregated score (values ranging between 0 and 2). For cases in which the microprotein could be targeted by more than eight sgRNAs, this aggregation score was used for sgRNA ranking. If various sgRNAs were less than four nucleotides apart, only the better ranked sgRNA was kept. Subsequently the best-ranking eight sgRNAs were included in the library. This resulted in a total of 50,136 sgRNAs targeting 11,776 unique microprotein candidates. Of note, since some sORFs displayed overlap, a number of sORFs with <2 (461) or >8 (1091) sgRNAs were retained, since these sgRNAs were included in the 2-8 sgRNAs targeting another ORF. Additionally, 1000 non-targeting negative control sgRNAs and 292 positive control sgRNA targeting ribosomal genes were included in the library.

#### Library cloning

The customised sORF library was synthesised as oligo nucleotides by Twist Bioscience **(Table S2)**. The array oligos were double-stranded and amplified via PCRs. The resulting PCR product included an A-U flip in the tracrRNA (74) 10-nucleotide random sequence labels (RSLs), and an i7 sequencing primer binding site (75) ggctttatatat**cttgtggaaaggacgaaacaccgnnnnnnnnnnnnnnnnnnnngtttaagagctagaaatagca agtttaaataaggct**agtccgttatcaacttgaaaaagtggcaccgagtcggtgctttttt<u>GATCGGAAGAGCACACG</u> <u>TCTGAACTCCAGTCAC</u>NNNNNNNNNNaagcttggcgtaactagatcttgagacaaa (bold: array oligo, n: sgRNA-sequence, N: RSL sequence, underlined: i7 sequence).

This construct was then cloned into the pLenti-Puro-AU-flip-3xBsmBI (Addgene #196709) (75) by Gibson assembly. After input sequencing to confirm library representation (same as gDNA sequencing described below but without PCR1), the plasmid library was packaged into the lentivirus utilising plasmids psPAX2 (a gift from Didier Trono, Addgene #12260) and pCMV-VSV-G (a gift from Bob Weinberg, Addgene #8454) in HEK-293T (ATCC). The lentivirus-containing supernatant was concentrated around 40-fold using Lenti-X concentrator (Takara), aliquoted and stored in liquid nitrogen.

#### Cas9 cell line generation

Cells stably expressing a codon optimized, WT SpCas9 flanked by two nuclear localization signals and coupled via a self-cleaving peptide to a blasticidin-S-deaminase-mTagBFP fusion protein (hereien called “A375 Cas9”, “HCT116 Cas9” or “K562 Cas9”) were generated by lentiviral transduction of the lenti.Cas9.BFP.Blast plasmid (Addgene #196714) and selected with 5 μg/ml blasticidin (Invivogen, ant-bl-10p). Subsequently, the cells were bulk sorted using a BD FACSAria Fusion (BD BioSciences) flow cytometer until a stable BFP-positive population was reached. Cas9 expression was additionally confirmed by Western blot.

#### CRISPR screening

The functional virus titre was estimated through assessment of cell survival rates after transduction of the respective cancer cell lines with different concentrations of virus and puromycin selection. For the screens, A375 Cas9, HCT116 Cas9 or K562 Cas9 cells (see “cell line generation”) were subjected to lentiviral transduction in duplicate at a MOI of around 0.3 and a coverage of around 1,000 cells per guide in presence of 2 µg/ml polybrene (Sigma-Aldrich, TR-1003-G). Two days after transduction, cells were selected with 2 μg/ml puromycin (VWR, CAYM13884) for five days and subsequently split into the different screening condition arms. For the essentiality screens, cells were grown for 21 days (A375 and HCT116 essentiality screens) or 28 days (K562 screens) after transduction and split every two to three days. A cell number of at least 60 million cells (around 1000 cells/sgRNA) per replicate was maintained at all times to ensure full library coverage. Cell pellets of 60 million cells were collected on day four and day 21 (A375, HCT116) or day seven and day 28 (K562) post– transduction for sequencing. For the 6-thioguanine screens, cells were treated with 30 μM 6-thioguanine (Sigma-Aldrich, A4882) in NaOH (stock solution 30mM 6-thioguanine in 1M NaOH) starting on day 10 post-transduction. The medium was subsequently changed every two to three days and cells were collected on day 28 post-transfection. Genomic DNA was extracted from the pellets with the QIAamp DNA Blood Maxi Kit (Qiagen). sgRNA and RSL sequences were amplified by three PCRs as described in (75) but with altered PCR2_FW and PCR3_fw primers. The amplicon was sequenced on Illumina NovaSeq, reading 20 cycles Read 1 with the CRISPRSEQ primer, 10 cycles index read i7 to read the RSL, and six cycles index read i5 for the sample barcode.

#### CRISPR screen analysis

The NGS data from the CRISPR screens was analysed using the MAGeCK software, version 0.5.8 (76) Log fold change (LFC) was calculated between sequencing reads of D4/D21 (A375, HCT116 essentiality), D7/D28 (K562 essentiality) or D28 control/D28 6-TG treatment (A375, K562 6-TG screens) with sORFs displaying a LFC ≤ -1 or LFC ≥ 1 considered to be hits. Full results can be found in **Table S3, S4.** For summary statistics, e.g. in Fig 1B,C,D, the sORFs were further deduplicated, retaining only unique sequences/genomic coordinates. To test for protein-coding (PC) overlap, we intersected the sORF coordinates with the CCDS database version 22 (77).

### Cell culture maintenance

A375, HCT116 and K562 parental cells were obtained from Sigma-Aldrich. A375 and HCT116 were grown in Gibco DMEM (Thermo Fisher Scientific, 10569010) supplemented with 10% FBS (Sigma-Aldrich, F7524). K562 cells were cultured in RPMI-1640 (Sigma-Aldrich, R8758) with 10% FBS (Sigma-Aldrich, F7524). During the CRISPR screening, the cells were maintained in 1x Pen-Strep (Sigma-Aldrich, P4333) at all times.

### Cell line generation

To generate A375 Cas9 cells (see “Cas9 cell line generation”) that additionally stably express GFP, we transduced the cells with lentivirus generated from pCDH_EF1_GFP_IRES_Puro. For viral packaging and production, the transfer plasmid was co-transfected with the envelope plasmid pMD2.G (Addgene #12259) and the second-generation packaging plasmid psPAX2 (Addgene #12260) into Hek293T cells (ATCC) using transIT®-LT1 (Mirus Bio). The growth media was replenished one day later. Viral particles were harvested 48- and 72-hours post-transfection and filtered through a 0.45 μm mixed cellulose esters syringe filter (Millipore). The first transduction was carried out on 40% confluent A375 Cas9 cells, seeded one day prior, by adding equivalent amounts of fresh DMEM and viral supernatant. 2 μg/ml polybrene (Sigma-Aldrich) was added to the growth media to boost efficiency of transduction. This was repeated with the second viral harvest and eight hours after the second transduction, cells were selected for five days in presence of 3 μg/ml puromycin (VWR, CAYM13884). A near 100% GFP-positive population was confirmed via microscopy (ZOE Fluorescent Cell Imager, BioRad) and flow cytometry (Navios flow cytometer, Beckman Coulter).

To generate microprotein rescue cells lines, the endogenous sORF nucleotide sequence was extracted, and every codon changed to the most common (using the codon efficiency table from (78)). ATG was used as the start codon and TAG TAA were used as two subsequent stop codons for each of the constructs. The corresponding oligos were ordered from Twist BioScience and cloned into an untagged (System BioSciences, PB533A-2), GFP-tagged (pPB_EF1_MCS-EGFP_IRES_Puro) or HA-tag backbone (pPB_EF1_MCS-HA_IRES_Puro) plasmid. Stable cell lines were generated using the PiggyBac Transposase system, in which A375 Cas9 cells were co-transfected with the rescue plasmids as well as the Super PiggyBac Transposase Expression Vector (System Bioscience Inc, PB200PA) at a 4:1 ratio utilising Lipofectamine™ LTX reagents (Thermo Fisher Scientific, 15338100) according to the manufacturer’s instructions. After 48 hours, cells were selected with 1-2 mg/ml G418 (Sigma-Aldrich, G8168) (untagged rescue plasmids) or 10 μg/ml puromycin (VWR, CAYM13884) (tagged rescue plasmids) for 5-7 days.

### Microscopy viability assay

Viability assays were carried out as three independent experiments with triplicate wells each (9 replicates total). A375 Cas9 cells were seeded at low confluency (2000 cells per well) into a 96-well plate (Falcon, 353219). One day after seeding, cells were transfected with the respective synthetic sgRNAs (Synthego) using Lipofectamine™ RNAiMAX transfection reagent (Thermo Fisher Scientific, 13778075) according to the manufacturers’ instructions. The medium was changed one day after transfection and live cells were imaged on an IN Cell Analyzer 2200 (GE Healthcare) at 4x magnification before the plate was returned to the incubator and cells were allowed to expand. Once they reached near confluency (day 4-6), cells were fixed and stained in a 4% formaldehyde (Thermo Fisher Scientific, 28908) and 2 μM Hoechst 33342 (Thermo Fisher Scientific, 62249) in PBS solution for 20 minutes, washed three times with PBS and then imaged in PBS with the IN Cell Analyzer 2200 (GE Healthcare) at 4x magnification. Cell numbers were counted via CellProfiler 3.1.9 (79). Briefly, cells were segmented on the basis of Cas9-BFP fluorescence (Day 1) or Hoechst fluorescence (Day 4-6), using the MaskImage function with customised masks, Gaussian filter smoothing, division illumination correction, speckle enhancement and adaptive three class thresholding with the Otsu method. Fold changes D4-6/Day 1 were normalised to the average triplicate TRAC fold change of each experiment. The average normalised fold change of the 9 replicates was then plotted in the figure and significance compared to TRAC control tested using a two-tailed unpaired student’s t-test.

### Knock-in generation

#### Plasmid generation

Knock-ins were performed using the PITCH method as previously described (Sakuma et al., 2016). Briefly sORF-specific template plasmids were created by AccuPrime Supermix PCR (Thermo Fisher Scientific, 12344040) using the pPB_EF1_MCS-EGFP_IRES_Puro as a PCR template and utilising primers with the respective knockin-overhangs (ordered from Sigma-Aldrich). Gel-extracted PCR products (Thermo Fisher Scientific, K0692) were then cloned into the MluI (Thermo Fisher Scientific, FD0564)-digested pCRIS-PITChv2-FBL plasmid (addgene #63672) using the In-Fusion HD cloning kit (Takara Bio, 639650). To create the sgRNA-plasmid backbone, PITCh sgRNA from pX330S-2-PITCh (Addgene #63670) was cut-out via Eco31I (Thermo Fisher Scientific, FD0293) and inserted into the pX330A-1x2 backbone (Addgene #58766) via a Golden gate assembly (T4 DNA ligase, NEB, M0202). sORF-specific sgRNAs were designed using the CRISPOR tool (56, 57), and ordered as DNA oligos (forward and reverse) from Thermo Fisher Scientific. The oligos (1:1 ratio) were phosphorylated and annealed in one step, using T4 PNK (Thermo Fisher Scientific, EK0032) according to the manufacturer’s recommendations but altering the incubation conditions (37°C for 20 minutes, 75°C for 10 minutes, 95°C for 5 minutes, 95°C - 25°C decrease at 6°C per minute). The thus-generated inserts were then ligated into the BpiI (Thermo Fisher Scientific, FD1014)-digested px330A-1x2_ PITCh_sgRNA plasmid using the T4 DNA ligase kit (Thermo Fisher Scientific, EL0016). For SNHG1_lnc_47, we did not find any suitable sgRNA that was U6-compatible. Thus, we ordered a synthetic sgRNA (Synthego) for this knock-in.

#### Transfection

1.6 Mio A375 parental cells were seeded. Two days after seeding, cells were double-transfected with sgRNA and template vector plasmids at a ratio of 2:1 using the Lipofectamine™ LTX reagents (Thermo Fisher Scientific, 15338100) and according to the manufacturer’s recommendation. Cells were expanded and bulk-sorted with the Sony SH8000 sorter two or three times to enrich for GFP+ cells (protein extracts of these polyclonal cells are shown in **Fig. 2A**), before being sorted as single cells into 96-well plates. Clones were trial-screened for positive PCR products by making mirror plates and subjecting one of the plates to cell lysis (50mM Tris-HCl pH 8, 1mM EDTA, 0.5% Tween-20, 50-80 μg/ml Proteinase K Merck-3115801001) at 37°C overnight. The resulting crude gDNA was transferred to a PCR plate, heated to 95°C for 10 min to inactivate the PK and performing genomic PCR with Phusion Mix (Thermo Fisher Scientific, F-548L) as described below but at 35 cycles and visualised with GelGreen (VWR, 730-1535) on a 1% agarose gel (run for 30 minutes at 135 V) using an ImageQuant™ LAS 500 (GE Lifesciences). Positive clones were then expanded from the remaining plate and genomic PCR and sequencing performed as outlined below.

For the SNHG1_lnc_47 knock-in, transfections of template (DNA) and sgRNA (RNA) were done in a sequential manner: Seeding was as above and two days after seeding, we transfected the template plasmid using Lipofectamine™ LTX reagents (Thermo Fisher Scientific, 15338100). The medium was changed and synthetic sgRNA transfected using Lipofectamine™ RNAiMAX transfection reagent (Thermo Fisher Scientific, 13778075) one day later. Expanding and sorting was done as above (though no GFP+ cells were observed).

#### Genomic PCR and sequencing

Genomic DNA was extracted with the GeneJET Genomic DNA Purification Kit (Thermo Scientific, K0721) according to the manufacturer’s recommendation. Genomic PCR primers were designed using the Primer-BLAST tool (80) and ordered from Thermo Fisher Scientific. Genomic PCR of 20 cycles was then performed with the respective primer pair on the extracted DNA utilising the Phusion Flash High-Fidelity PCR Master Mix (Thermo Fisher Scientific, F-548L) as described by the manufacturer. PCR products were visualised on a 1% agarose gel (run for 30 minutes at 135 V) with GelGreen (VWR, 730-1535) using an ImageQuant™ LAS 500 (GE Lifesciences), excised and DNA extracted using the GeneJet Gel extraction kit (Thermo Fisher Scientific, K0692). PCR products were subsequently sequenced by using the primers that were also utilized in the genomic PCR and Eurofins Genomics sequencing service.

### Western blotting

Cells were pelleted at 300 x g for 5 minutes, pellets washed once with PBS (Sigma-Aldrich, D8537), spun again and then resuspended in ice-cold RIPA-SDS buffer (50 mM Tris-HCl pH 7.6, 150 mM NaCl, 0.25 % sodium deoxycholate, 1mM EDTA, 1% NP-40, 0.1% SDS), supplemented with Complete Protease inhibitor (Sigma-Aldrich, 5056489001). To gain the soluble protein fraction, protein extracts were centrifuged at maximum speed for ten min at 4°C in a table-top centrifuge and the supernatant was extracted. SDS buffer (final concentration: 62.5 mM Tris pH 6.8, 2% SDS, 10% glycerol, 0.1M DTT, 0.01% bromophenol blue) was added, the sample boiled for five minutes at 95 °C and size-separated by SDS-PAGE using a 4–20% Mini-PROTEAN® TGX™ Gel (Biorad). Proteins were transferred onto a nitrocellulose membrane (Biorad, 1704270) and Ponceau staining (Sigma-Aldrich, P7170) was performed. Blocking and antibody incubations were done in 4% TBS-T under mild shaking. The membrane was blocked for one hour at room temperature, antibody incubations were done overnight at 4 °C and secondary antibody incubations were done for one hour at room temperature. Proteins were visualised using Immobilon Classico Western HRP substrate (Sigma-Aldrich, WBLUC0100). Primary antibody dilutions are as follow: anti-GFP 1:5000 (B-2, mouse monoclonal, Santa Cruz sc-9996), anti-β-Actin 1:4000 (13E5, Cell Signaling 4970, rabbit monoclonal). Secondary antibodies as follows: anti-mouse 1:10,000 (BioRad 1721011), anti-rabbit 1:10,000 (BioRad 1721019).

### Co-immunoprecipitation experiments

#### Sample preparation

For each of the conditions, we prepared two biological replicates. For each replicate, a confluent T75 flask of cells was harvested using TrypLE Express (Thermo Fisher Scientific, 12605010) and pelleted for five minutes at 300 x g. Pellets were washed with PBS (Sigma-Aldrich, D8537), the suspension centrifuged for five minutes at 300 x g again and the resulting pellet flash frozen in dry ice and Methanol (VWR, 20847.307). Pellets were thawed on ice, resuspended in ice-cold RIPA buffer (50 mM Tris-HCl pH 7.6, 150 mM NaCl, 0.25 % sodium deoxycholate, 1mM EDTA, 1% NP-40) supplemented with Complete Protease inhibitor (Sigma-Aldrich, 5056489001) and incubated for five minutes on ice. The resulting protein extracts were centrifuged at maximum speed on a table-top centrifuge for 15 minutes at 4 °C and the supernatant (soluble fraction) incubated with 25 μl GFP-trap magnetic beads (ChromoTek, gtma) for four hours at 4 °C under constant rotation. The protein-bound beads were then washed once with RIPA buffer, twice with PBS or wash buffer (0.5 % NP-40, 0.1 mM EDTA, 20 mM Tris–HCl pH 7.4, 500 mM NaCl) and once with ddH20. Bound proteins were subsequently eluted twice for five minutes with 15 μl 1% acetic acid. Protein eluates were reduced and alkylated in the same step with 5 mM TCEP (Thermo Fisher Scientific, PG82080) and 20 mM chloroacetamide (Sigma-Aldrich, C0267) in 250 mM Tris buffer pH 8 for 30 minutes at room temperature. Subsequently, Sera-Mag magnetic bead mix (1:1 ratio of Sigma-Aldrich, GE45152105050250 and GE65152105050250) was added in a 25:1 protein to bead volume ratio and proteins were bound onto the beads by adding an equal volume (protein plus beads) of absolute ethanol. After a five-minute incubation, the supernatant was removed and beads were washed three times with 80% ethanol. To perform on-bead digestion, protein-bound beads were resuspended in 50mM TEAB pH 8 (Sigma-Aldrich, T7408) containing 1 μg trypsin (Thermo Fisher Scientific, 90057) and incubated overnight gently shaking at 37 °C. Afterwards, beads were allowed to settle on the magnetic rack and the supernatant was taken and kept as the first protein eluate. Subsequently, another elution was performed by adding 50 μl 2% acetonitrile (Sigma-Aldrich, 34851) in 50 mM TEAB. The second elution was combined with the first one and the eluate was dried in a SpeedVac.

#### LC-MS/MS

LC-MS/MS analysis was performed by using an EASY-nLC 1200 (Thermo Fisher Scientific) coupled to an Exploris 480 mass spectrometer (Thermo Fisher Scientific). Separation of peptides was performed on 20 cm frit-less silica emitters (CoAnn Technologies, 0.75 µm inner diameter), packed in-house with reversed-phase ReproSil-Pur C18 AQ 1.9 µm resin (Dr. Maisch). The column was constantly kept at 50 °C. Peptides were eluted in 115 min applying a segmented linear gradient of 0 % to 98 % solvent B (solvent A 0 % ACN, 0.1 % FA; solvent B 80 % ACN, 0.1 % FA) at a flow-rate of 300 nL/min. Mass spectra were acquired in data-dependent acquisition mode. MS1 scans were acquired at an Orbitrap Resolution of 120,000 with a Scan Range (m/z) of 380-1500, a maximum injection time of 100 ms and a Normalised AGC Target of 300 %. For fragmentation only precursors with charge states 2-6 were considered. Up to 20 Dependent Scans were taken. For dynamic exclusion, the exclusion duration was set to 40 sec and a mass tolerance of +/- 10 ppm. The Isolation Window was set to 1.6 m/z with no offset. A normalised collision energy of 30 was used. MS2 scans were taken at an Orbitrap Resolution of 15,000, with a fixed First Mass (m/z) = 120. Maximum injection time was 22 ms and the normalised AGC Target 50 %.

#### Data analysis

The raw data was analysed using MaxQuant Version 1.6.3.4 (81) and searched against the Uniprot protein database (71) (Human all 2017/11), as well as our microprotein library, a list of GFP-fused bait candidates and the CRAPome database (82) for contaminants. MQ default settings were used, besides the following adjustments: Match between runs and LFQ intensity reporting were activated and MS/MS identification for both peptides in a pairwise LFQ intensity comparison was not required. The resulting proteinGroups file was analysed using the DEP package 1.22.0 (83). Common contaminants were filtered out and imputation performed according to the “min” method. No DEP normalisation was carried out. For a protein to be considered a hit, we required its presence in at least three out of four replicates for the endogenously expressed CENPBD2P_p_13 knock-in CO-IP or 6 out of 6 replicates for the overexpressed fusion protein CO-IP. Additional requirements were a LFC enrichment of ≥ 1 (CENPBD2P_p_1 knock-in) or ≥ 1.5 (all others) over the parental control and an adjusted DEP p-value of ≤ 0.05. The remaining candidates were intersected to test for overlap, and any protein present in more than one pulldown was excluded from further analysis. Volcano plots of the resulting data were generated using R (https://www.R-project.org/, 4.2.1) and Rstudio (http://www.rstudio.com, 2022.07.1) and further processed with Adobe Illustrator. MS/MS spectra plots shown for the CENPBD2P_p_13 knock-in were generated via MaxQuant Version 1.6.3.4 (81).

### Genomic location and conservation alignments

The genomic location (GRCh38 assembly) and phylogeny tree were outputted from the UCSC Genome Browser (84). Genomic alignment was performed using the Cons 30 Primates track and alignment subsequently manually confirmed. The respective genomic sequences were then translated using the Expasy translation tool (85) and the resulting amino acid sequence pasted into the EMBL-EBI Clustal Omega tool (86) and from there transferred into JalView (87) for colouring.

### Confocal microscopy

For live cell imaging 10,000 cells/well were seeded in a 18-well glass bottom plate (Ibidi, 81817). 24 hours after seeding, cells were washed once with PBS and subsequently incubated with prewarmed live cell imaging solution (Thermo Fisher Scientific, A14291DJ) containing 2 μM Hoechst 33342 (Life Technologies, 62249) for 20 minutes at 37°C. The staining solution was removed, cells were washed twice with PBS and then z-stack imaged in fresh pre-warmed live cell imaging solution. For fixed cell imaging, cells were seeded as above. 24 hours after seeding, wells were washed twice with PBS, fixed and stained using 4% formaldehyde (Thermo Fisher Scientific, 28908) and 2 μM Hoechst 33342 (Thermo Fisher Scientific, 62249) in PBS for fifteen minutes. Three PBS washes were performed and cells were subsequently z-stack imaged in PBS. Imaging was performed using a Zeiss LSM800-Airy Scan laser scanning microscope with a 60x oil immersion objective. Shown is a single plane from the z-stack image. Images were analysed using Fiji/ImageJ and annotated in Adobe Illustrator.

### Motif searches

Motif searches were performed using the ELM tool (88).

### Gene ontology analysis

Gene ontology analysis was performed using gprofiler version 20230223 (89), MetaScape version v3.5.20230501 (90) and String database version 11.5 tools (91) with default parameters.

### Venn Diagrams

Venn Diagrams were made with InterActiVenn (92).

### Rescue assays

The generated rescue cell lines and A375 Cas9 or A375 Cas9 GFP cells were seeded at a 1:1 (80,000 cells/well total) ratio into a 24-well plate. One day after seeding, cells were transfected with synthetic sgRNAs (Synthego) utilizing the Lipofectamine™ RNAiMAX transfection reagent (Thermo Fisher Scientific, 13778075) according to the manufacturer’s instructions. One day after transfection, cells were transferred into 12-well plates and grown for another seven days. Cells were split on days 1, 4, 6 and harvested on day 8 post-transfection. A small fraction of cells was separated for flow cytometry analysis on each splitting day and all cells were harvested on day 8 post-transfection. For all flow cytometry analysis, cells were harvested, pelleted for five minutes at 300 x g, cell pellets resuspended in 5% FBS (Sigma-Aldrich, F7524) in PBS (Sigma-Aldrich, D8537) and then analysed on a Navios flow cytometer (Beckman Coulter). Flow cytometry data analysis was performed with FlowJo™ v10.8 Software (BD Life Sciences), assessing single cell GFP+ and GFP-fractions.

### RNA-sequencing

#### Sample preparation

80,000 A375 Cas9 cells per well were seeded in a 24-well plate and transfected with the respective synthetic sgRNA (Synthego) in triplicate wells one day after seeding using Lipofectamine™ RNAiMAX (Thermo Fisher Scientific, 13778075) according to the manufacturer’s protocol. Cells were transferred into a 6-well plate one day post-transfection and expanded until three days post-transfection. One million cells were then harvested by trypsinization (TrypLE Express, Thermo Fisher Scientific, 12605010) and pelleted by centrifugation at 500 x g for three minutes. Pellets were washed once with PBS (Sigma-Aldrich, D8537), flash frozen and stored at -80C until use. The pellet was resuspended in 350ul RLT buffer (Qiagen, 74106) supplemented with 1% beta-mercaptoethanol. The lysate was homogenised using the Qiashredder column (Qiagen, 79654) and RNA extracted utilising the RNeasy Plus Mini Kit (Qiagen, 74136) according to manufacturer’s instructions. RNA concentration was measured with the Qubit RNA HS assay kit (Life Technologies, Q32852) according to manufacturer’s instructions and methanol flash frozen. RNA-sequencing service was provided through BGI services (www.bgi.com), performing strand-specific RNA-seq with poly(A) selection (DNBseq Eukaryotic Transcriptome De novo Sequencing).

#### Analysis

RNA-seq FASTQ files were processed using the nf-core RNA-seq pipeline version 3.5 (https://nf-co.re/rnaseq/3.5) with star_rsem parameters (93, 94), hg38 as reference and RefSeq as gene annotation. The resulting counts were processed using the DeSeq2 software (95) with default parameters to calculate corresponding LFC and significance values. TPM values displayed as calculated by the pipeline. For a gene to qualify as a hit, we required an average LFC of ≥ 0.5 or ≤ –0.5 between microprotein sgRNA and TRAC sgRNA treated A375 Cas9 cells, a DeSeq2 adjusted p-value of ≤ 0.05 and the average gene expression variability of the two TRAC conditions (microprotein-GFP vs. parental cells) to not exceed 10%. Subsequently we scored rescued hits, by testing for genes that additionally displayed a LFC difference of ≥ 0.5 between the microprotein knockout conditions in microprotein-GFP and parental cell lines.

#### Ribo-seq analysis

Published Ribo-seq data (96) (GSE143263) was processed through the same pipeline as RNA-seq above.

### Immunofluorescent microscopy

10,000 cells/well were seeded in a 96-well glass bottom plate (VWR, GREI655891_16). 24 hours after seeding, the cells were washed once with PBS (Sigma-Aldrich, D8537), fixed with 4% formaldehyde (Thermo Fisher Scientific, 28908) for fifteen minutes at room temperature and subsequently washed twice time with PBS. Cells were permeabilized using 0.1% Triton X-100 (Sigma-Aldrich, T9284) in PBS (Sigma-Aldrich, D8537) for 15 minutes and washed twice with TBS-T. The samples were blocked for one hour at room temperature with 0.1% BSA in 0.05% Triton X-100 (Sigma-Aldrich, T9284-100ML) in TBS-T. For staining, primary antibodies were diluted in the blocking solution, and incubated with the sample for one hour at room temperature or overnight at 4°C. Subsequently, three washes with TBS-T were performed and the samples were incubated with secondary antibody diluted in blocking solution, for one hour at room temperature, followed by three washes with TBS-T. To stain nuclei, the cells were then incubated with 1μg/ml DAPI (Sigma-Aldrich, D9542) in PBS for five minutes and washed twice with PBS. The sample was imaged in PBS on a Nikon Eclipse Ti2 inverted widefield microscope. Primary and secondary antibody dilutions as follows: anti-HA 1:500 (F-7, mouse monoclonal, Santa-Cruz sc-7392), anti-mouse Alexa 647 1:1000 (Thermo Fisher Scientific, A-21236).

### Synthetic sgRNAs

All Synthetic sgRNAs were ordered from Synthego.

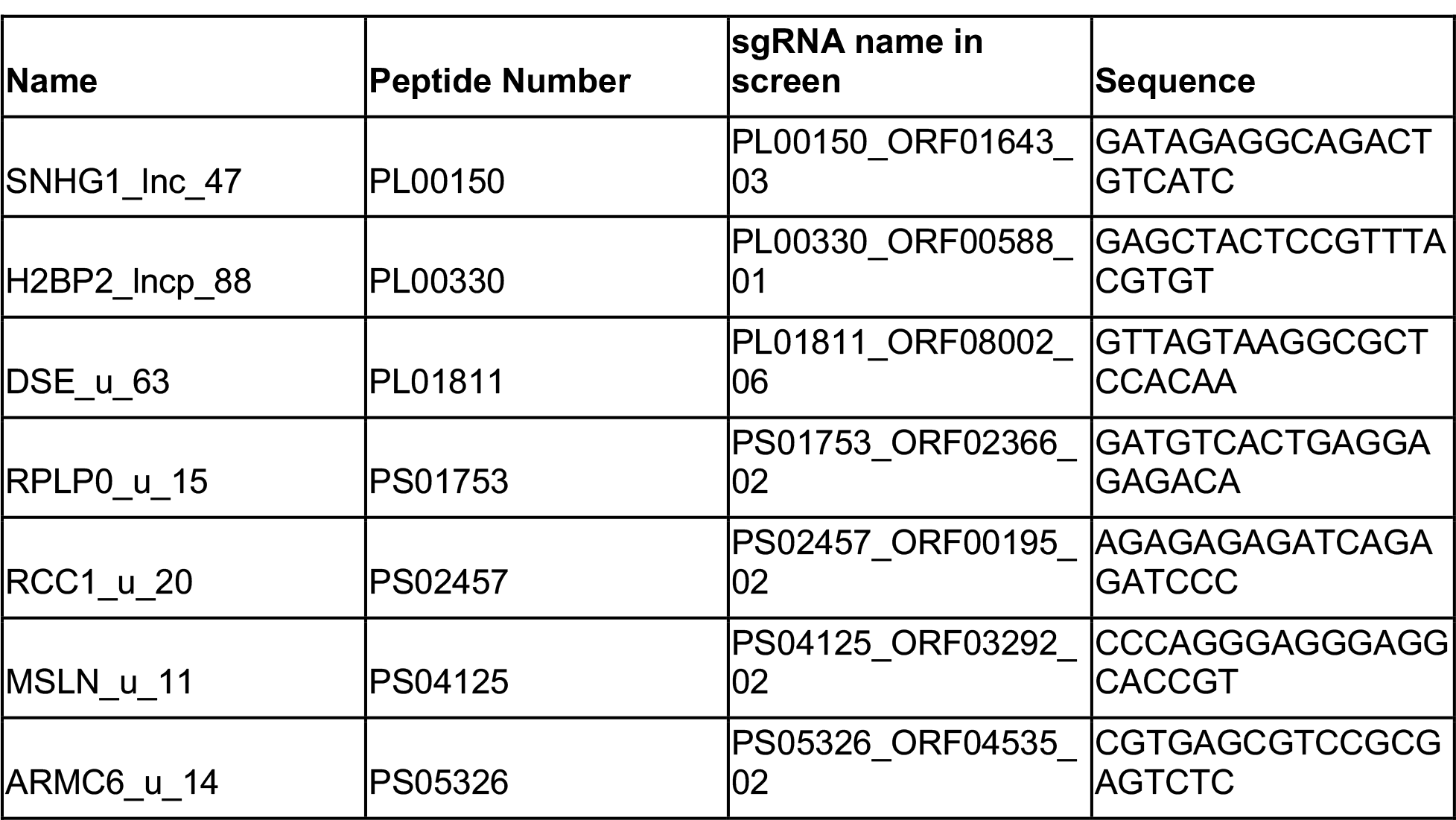
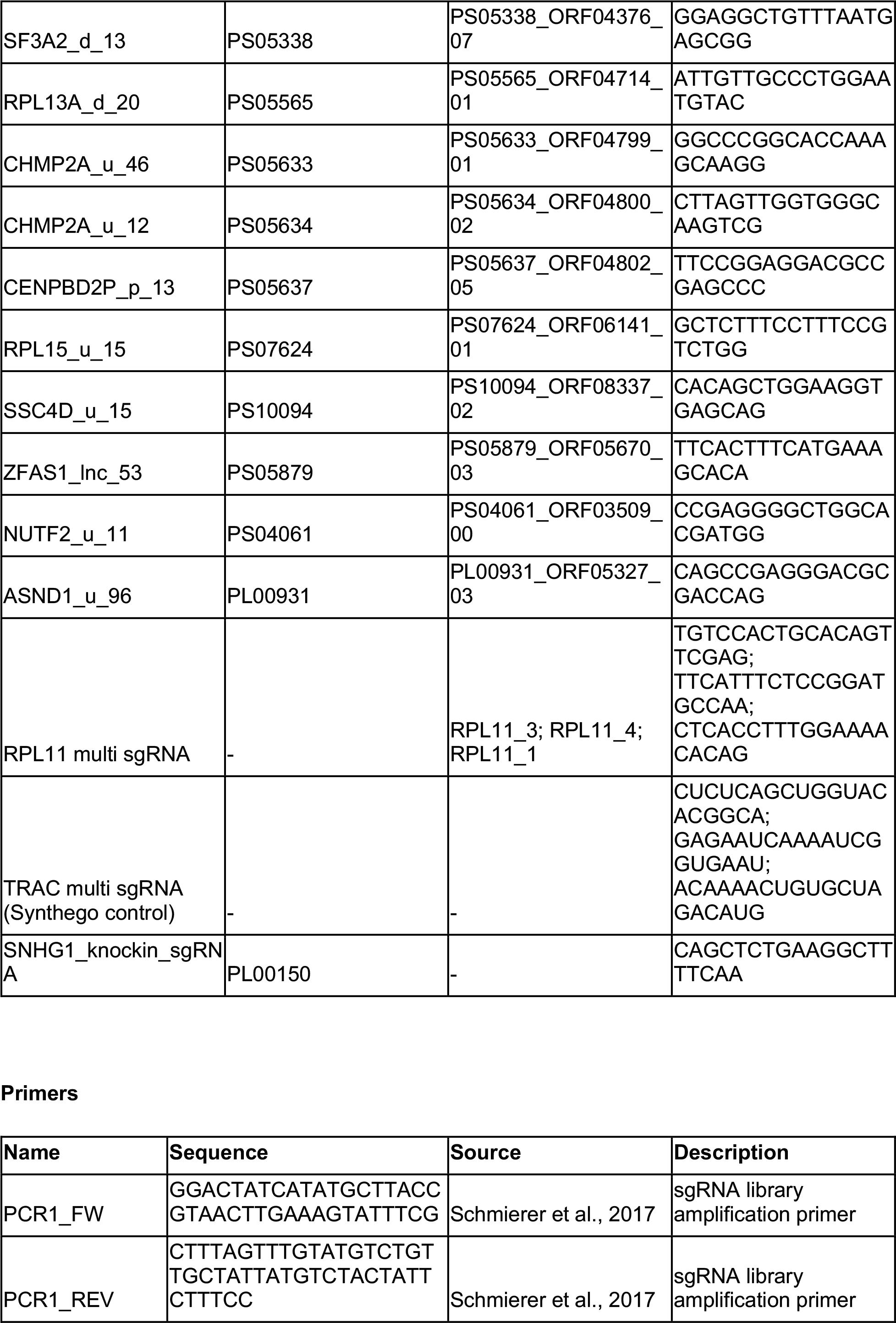
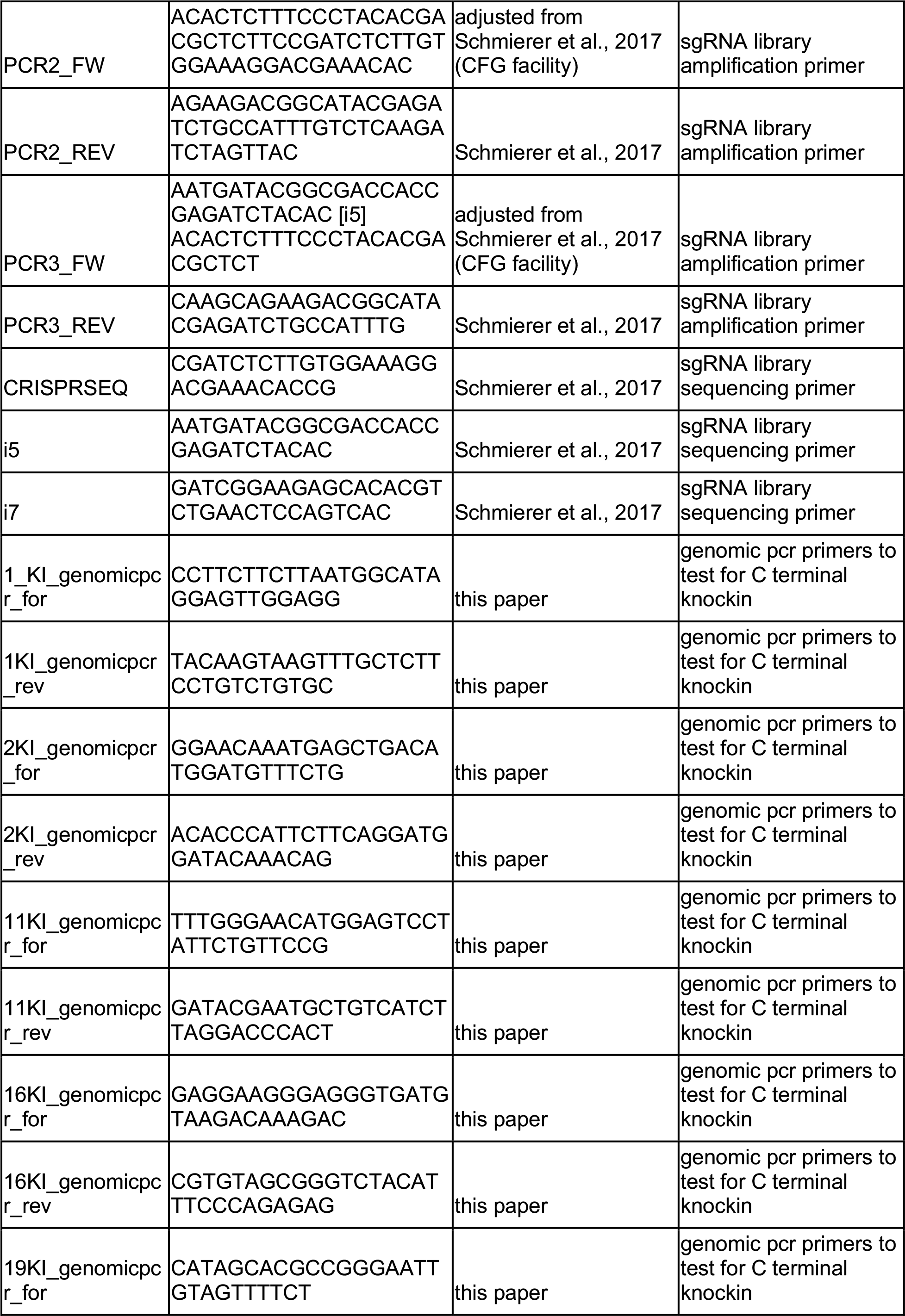
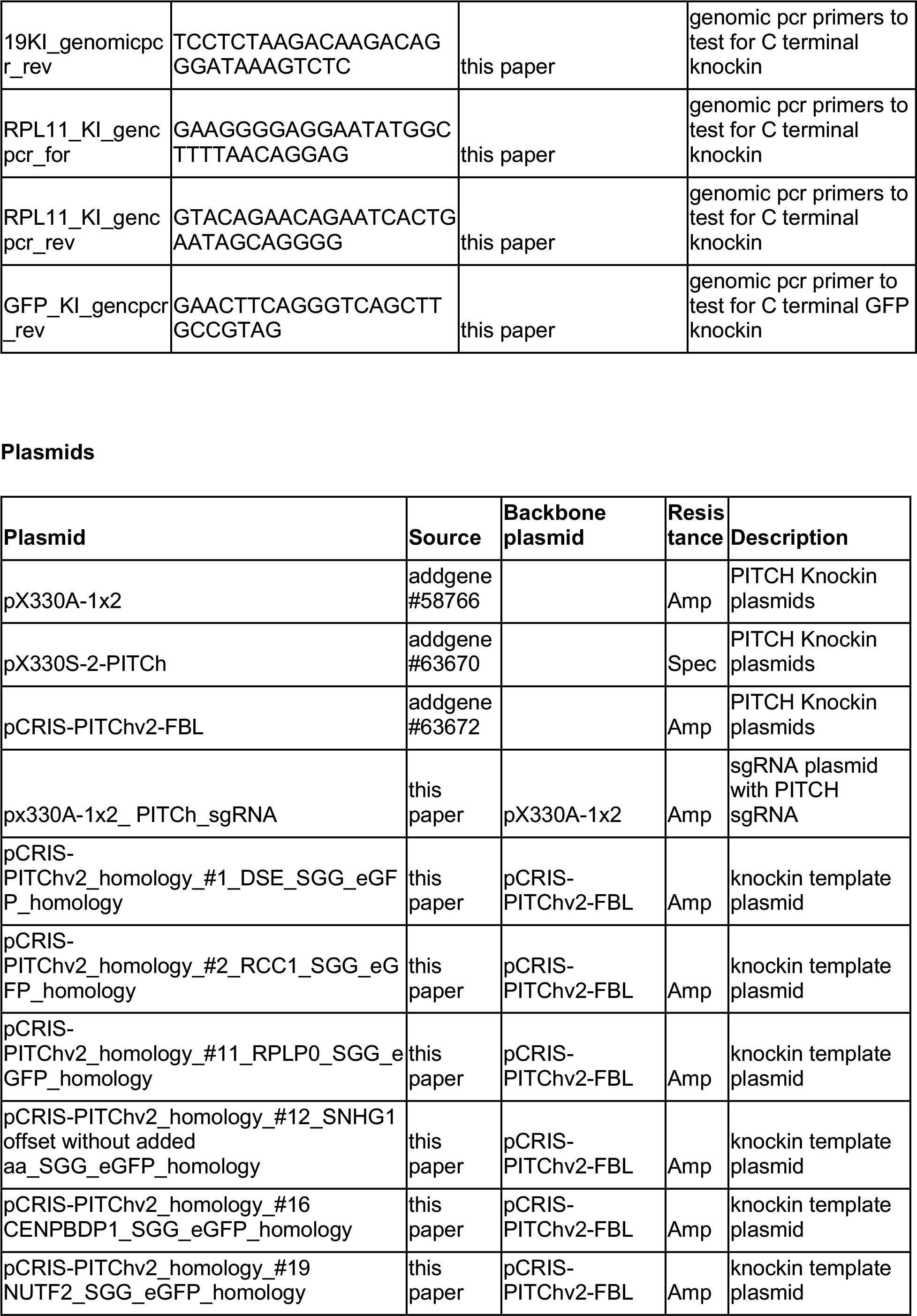
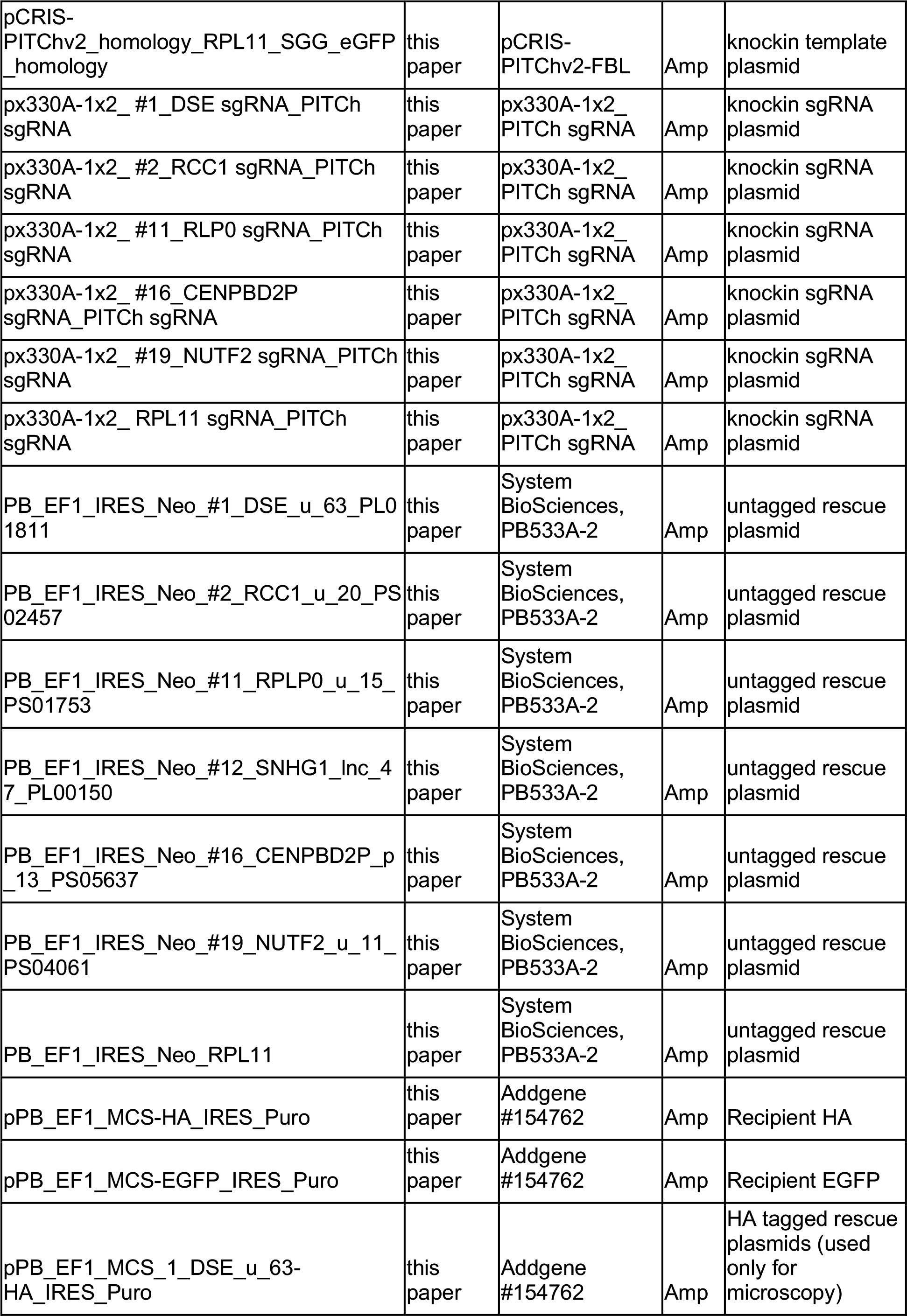
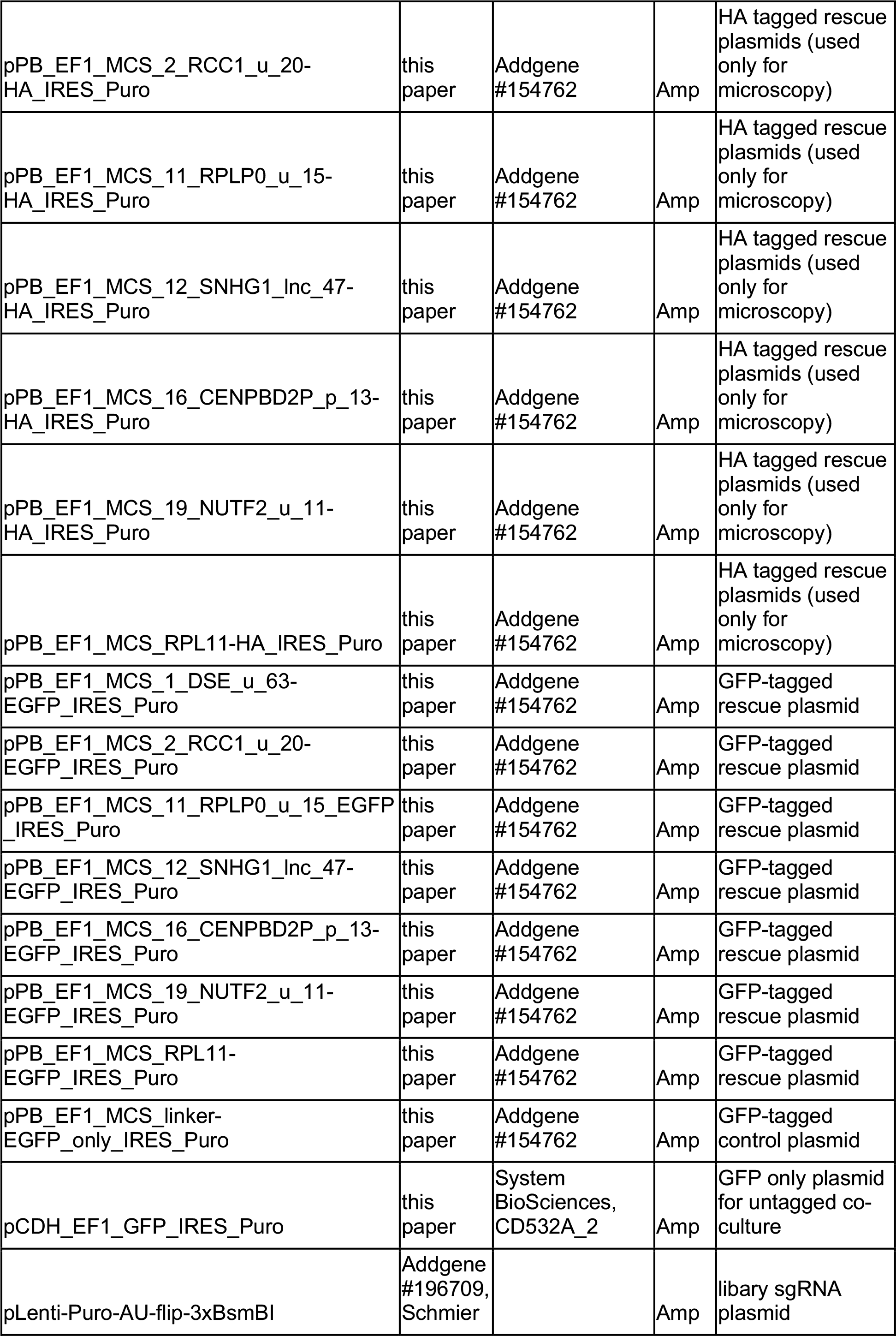
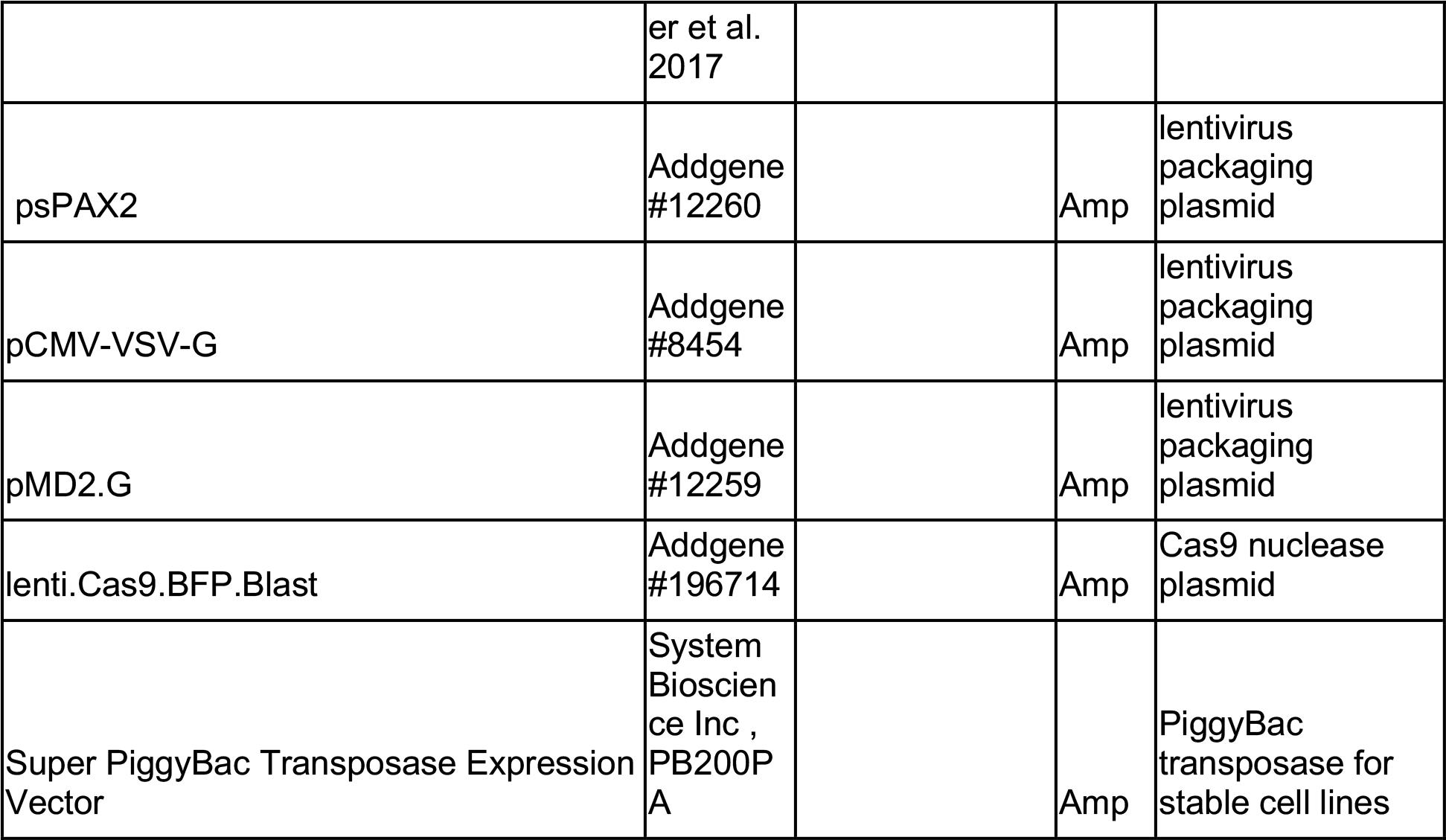

