## Supplementary Information for "A large-scale sORF screen identifies putative microproteins and provides insights into their interaction partners, localisation and function"

Supplementary Figures S1-S10
Supplementary Tables 1-4

### Supplementary Figure 1

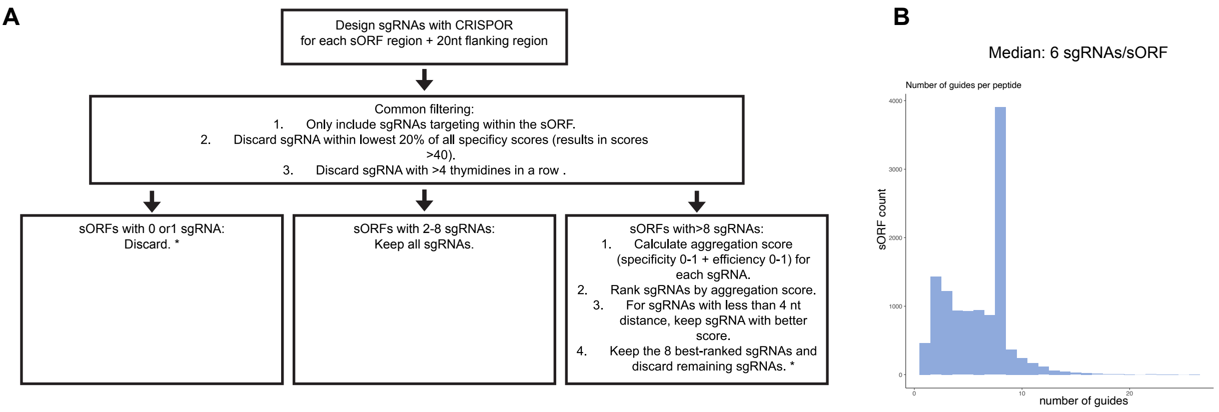


Supplementary Figure 1: sgRNA design for sORF CRISPR screening. A) Flow chart for sgRNA generation. * Of note, even though we filtered out sgRNAs that are single targeting guides or are not ranked amongst the top 8 sgRNAs for a respective sORF, these sgRNAs/sORFs might still be included due to overlap with other sORFs in which these sgRNAs score highly. This resulted in a small number of sORFs targeted by a single guide or by >8 sgRNAs (see also Supp Fig 1 B) B) Barplot showing number of sgRNAs/ORF.

### Supplementary Figure 2

**A**

| | A375 | HCT116 | K562 |
|---|---|---|---|
| 10% FBS | X | X | X |
| 1% FBS | | X | X |
| 0% FBS | | X | |
| 6-TG | X | | X |

**B**

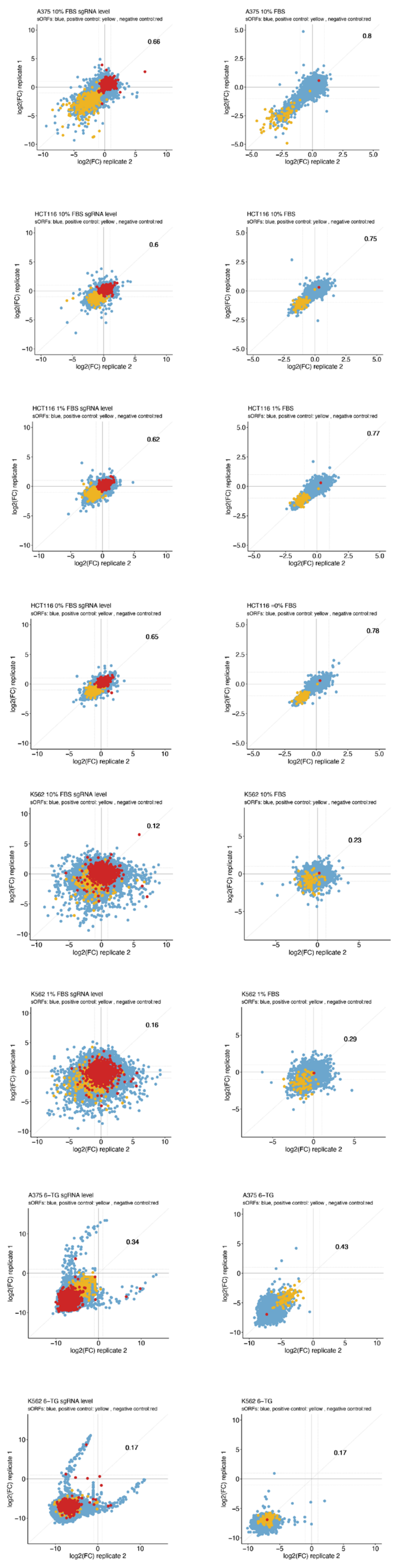


**C**

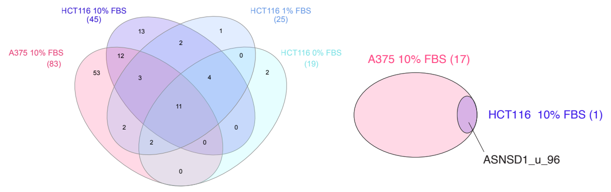


**D**

**E**

**F**

**G**

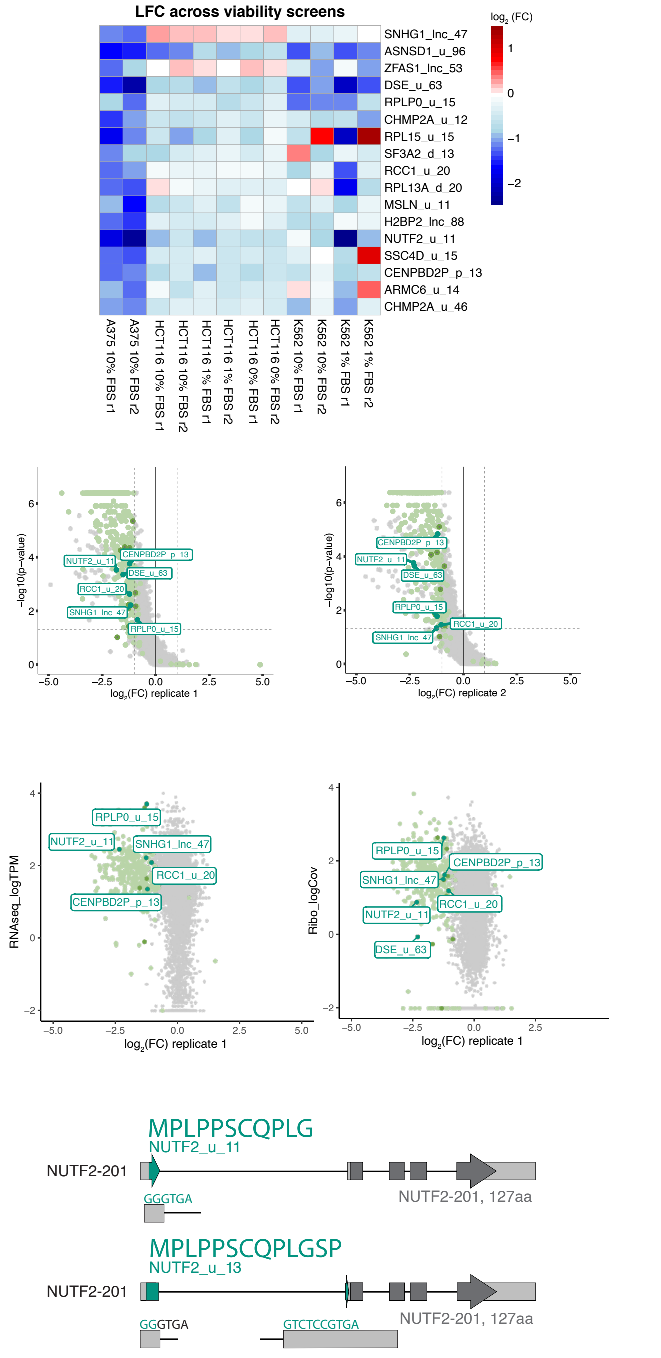


Supplementary Figure 2: Results of the sORF-specific screen in all cell lines. A) Table summarizing all screening condition, marked screens displayed good dynamic range and correlation between replicates. B) Scatterplots showing results of all carried out screens, and depicting log2 fold change (LFC) for individual sgRNAs (left panels) or median LFC of all sgRNAs in the two replicates (right panels). positive controls (sgRNAs targeting ribosomal genes): yellow; negative controls (non-targeting sgRNAs): red. correlation coefficients are indicated in the right corner. C) Venn diagram illustrating all hits (left) or the 17 candidates (right) in the A375 and HCT116 full serum screens. D) Heatmap showing median LFC of all sgRNAs targeting each of the 17 candidate ORF in all of the viability screening conditions. E) Volcano plots illustrating LFC and statistical significance (MAGeCK negative p-value) for each sORF in a given replicate of the 10% A375 screen. non-hits: grey, hits: light green, candidates: green, Top hits: dark green. F) Scatter plot showing RNA-seq TPM transcript-level expression (left) and average RiboSeq read coverage of the sORF (right) against LFC of A375 screen. non-hits: grey, hits: light green, candidates: green, Top hits: dark green. G) Schematic of originally described NUTF2_u_11 sORF and an alternative version in the NUTF2-201 transcript UTR. dark green arrow: sORF, grey arrow: canonical ORF, grey bars: exons, lines: introns.

#### Supplementary Figure 3 (related to Fig.2)

A

| Knock -in construct | Issue encountered |
|---|---|
| DSE_u_63 | Off-target (RPS5) integration and/or frameshifting indel |
| RCC1_u_20 | Off-target integration and/or frameshifting indel |
| RPLP0_u_15 | 28-nt insertion in all clones |
| SNHG1_lnc_47 | Lack of U6-compatible sgRNA, lack of GFP signal after knock-in with Synthego sgRNA+ plasmid template |
| CENPBD2P_p_13 | Knock-in successful |
| NUTF2_u_11 | Lack of PCR product |

B

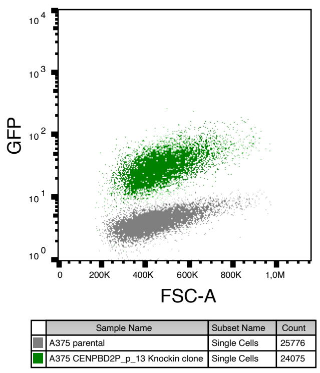


C

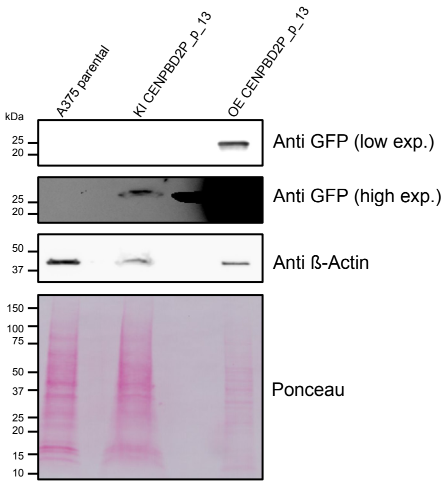


E

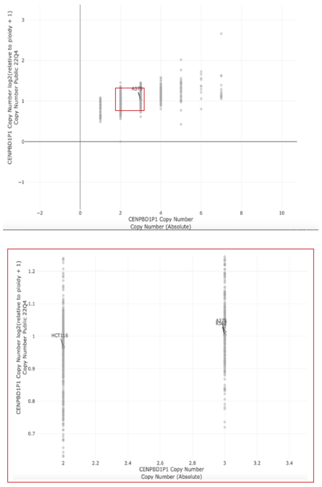

D

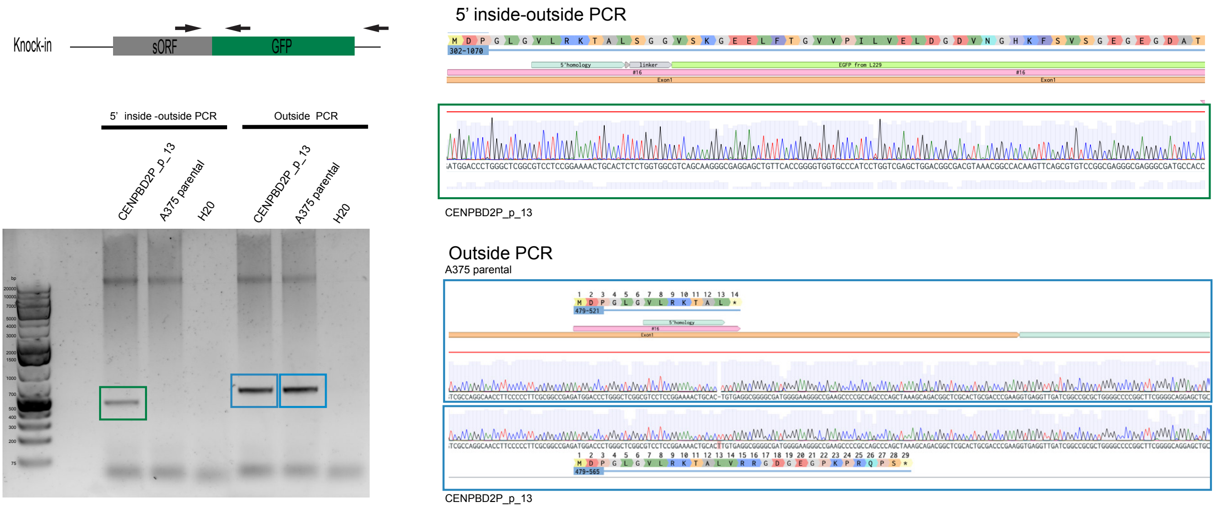


Supplementary Figure 3: Validation of CENPBD2P_p_13 is translation. A) Table summarizing issues in validation processes for each of the putative microprotein knock-ins. B) Scatterplot of flow cytometry GFP signal for CENPBD2P_p_13-GFP A375 knockin clone and A375 parental cells. C) Western blot showing CENPBD2P_p_13-GFP knockin clone (KI), CENPBD2P_p_13-GFP stable rescue cell line (OE) and A375 parental cells. D) Agarose gel (left panel) and sequencing results (right panel) of genomic PCR products with primers either binding outside of the integration region (”outside PCR”, blue), or a forward primer binding outside and reverse primer binding inside of the integration region (”inside-outside PCR”, green) in the CENPBD2P_p_13-GFP knockin clone and A375 parental cells. Only one band instead of two bands is visible in the knockin clone outside PCR, likely due to bias towards the much smaller non-integration PCR product.  E) Output of DepMap copy number for CENPBD2P locus with A375, HCT116 and K562 cell lines marked.

### Supplementary Figure 4 (related to Fig.2)

A

| Elm Name | Instances (Matched Sequence) | Positions | View in Jmol | Elm Description | Cell Compartment | Pattern | PHI-Blast Instance Mapping | Structural Filter Info | Probability |
|---|---|---|---|---|---|---|---|---|---|
| CLV_NRD_NRD_1 | LRK | 8-10 [A] | - | N-Arg dibasic convertase (NRD/Nardilysin) cleavage site (X-\|-R-K or R-\|-R-X). | extracellular, Golgi apparatus, cell surface | (.RK)\|(RR[^KR]) | - | - | 7.465e-03 |
| DEG_Nend_UBRbox_2 | MDP | 1-3 [A] | - | N-terminal motif that initiates protein degradation by binding to the UBR-box of N-recognins. This N-degron variant comprises N-terminal Asp or Glu as destabilizing residue. | cytosol | ^M{0,1}([ED]). | - | - | 2.537e-04 |
| LIG_PDZ_Class_1 | LRKTAL | 8-13 [A] | - | The C-terminal class 1 PDZ-binding motif is classically represented by a pattern like (ST)X(VIL)* | cytosol, internal side of plasma membrane | ...[ST].[ACVILF]$ | - | - | 7.255e-05 |
| MOD_NEK2_1 | LRKTAL | 8-13 [A] | - | NEK2 phosphorylation motif with preferred Phe, Leu or Met in the -3 position to compensate for less favorable residues in the +1 and +2 position. | centrosome, Ndc80 complex, condensed nuclear chromosome outer kinetochore, cytosol, nucleus | [FLM][^P][^P]([ST])[^DEP][^DE] | - | - | 9.798e-03 |

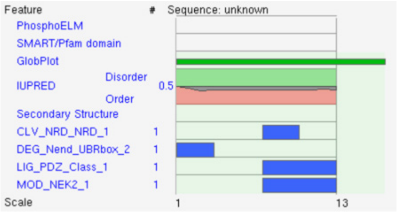


B

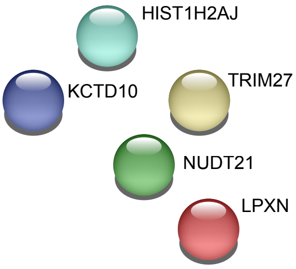


C

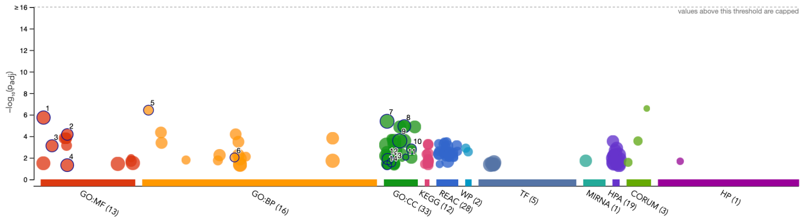


| ID | Source | Term ID | Term Name | Padj (query_1) |
|---|---|---|---|---|
| 1 | GO:MF | GO:0003723 | RNA binding | $1.823\times10^{-6}$ |
| 2 | GO:MF | GO:0017111 | ribonucleoside triphosphate phosphatase ac... | $6.706\times10^{-5}$ |
| 3 | GO:MF | GO:0005198 | structural molecule activity | $7.611\times10^{-4}$ |
| 4 | GO:MF | GO:0017076 | purine nucleotide binding | $4.817\times10^{-2}$ |
| 5 | GO:BP | GO:0002181 | cytoplasmic translation | $3.760\times10^{-7}$ |
| 6 | GO:BP | GO:0042776 | proton motive force-driven mitochondrial AT... | $9.221\times10^{-3}$ |
| 7 | GO:CC | GO:0005829 | cytosol | $4.041\times10^{-6}$ |
| 8 | GO:CC | GO:0070062 | extracellular exosome | $1.139\times10^{-5}$ |
| 9 | GO:CC | GO:0043233 | organelle lumen | $2.642\times10^{-4}$ |
| 10 | GO:CC | GO:0098800 | inner mitochondrial membrane protein compl... | $1.488\times10^{-3}$ |
| 11 | GO:CC | GO:0070937 | CRD-mediated mRNA stability complex | $8.671\times10^{-3}$ |
| 12 | GO:CC | GO:0005925 | focal adhesion | $1.293\times10^{-2}$ |
| 13 | GO:CC | GO:0030120 | vesicle coat | $2.725\times10^{-2}$ |
| 14 | GO:CC | GO:0005844 | polysome | $4.928\times10^{-2}$ |

Molecular Function (Gene Ontology)

| GO-term | description | count in network | strength | false discovery rate |
|---|---|---|---|---|
| GO:0003735 | Structural constituent of ribosome | 5 of 159 | 1.21 | 0.0083 |
| GO:0005198 | Structural molecule activity | 9 of 635 | 0.86 | 0.0019 |
| GO:0003723 | RNA binding | 21 of 1649 | 0.82 | 6.90e-10 |
| GO:0003676 | Nucleic acid binding | 22 of 3947 | 0.46 | 0.00031 |
| GO:1901363 | Heterocyclic compound binding | 27 of 5831 | 0.38 | 0.00031 |

(more ...)

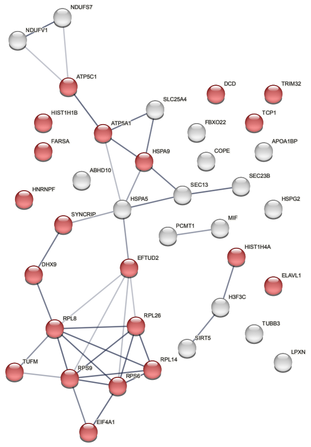


Supplementary figure 4: Further characterization of CENPBD2P_p_13 A) Output of ELM motif search. B) String database output for interaction partners identified in the harsh wash CENPBD2P_p_13 knockin Co-IP, gprofiler GO analsis yielded no results. C) String database (right panel) and gprofiler GO output (left panels) for interaction partners identified in the PBS wash CENPBD2P_p_13 knockin Co-IP.

### Supplementary Figure 5 (related to Fig.3)

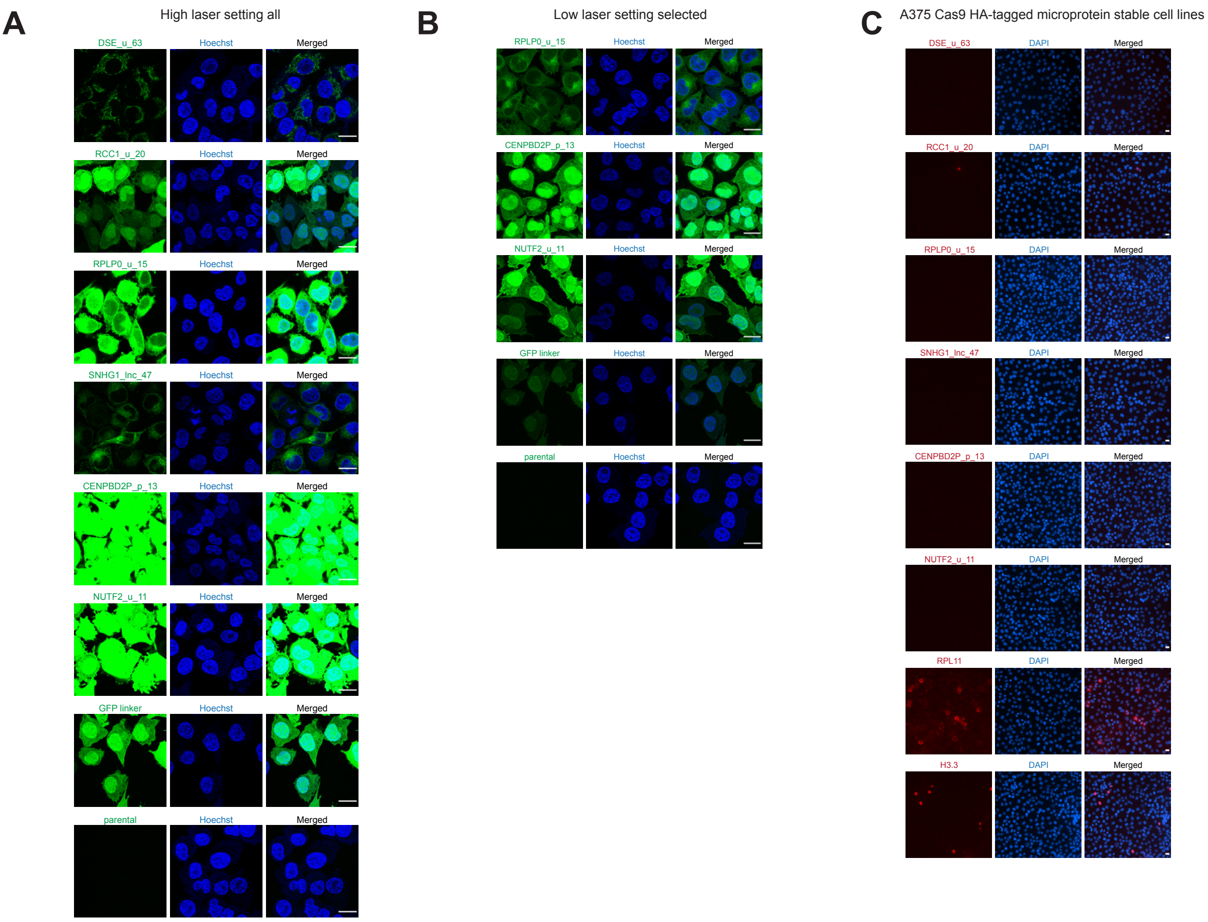


Supplementary figure 5: Microscopy of putative microproteins (A-B). Fixed confocal microscopy images of A375 Cas9 parental cells, cells stably expressing the respective codon-altered GFP-fused putative microproteins, or cells expressing the GFP linker only, acquired with higher laser setting (A) or lower laser setting (B). For figure 3, DSE_u_63, RCC1_u_20 and SNHG1_lnc_47 were acquired with the higher laser setting and RPLP0_u_15 and NUTF2_u_11 were acquired with the lower laser setting. Laser and Fiji settings were kept the same within each of the panels (A and B). green: GFP, blue: HOECHST 33342. Shown is a single plane from a z-stack. Scale bar represents 20 µm. C) Fixed widefield microscopy images of A375 Cas9 parental cells stably transfected with codon-altered HA-fused putative microprotein constructs or controls. red: HA-tag (Alexa 647), blue: DAPI. Scale bar represents 20 µm.

#### Supplementary Figure 6 (related to Fig.3)

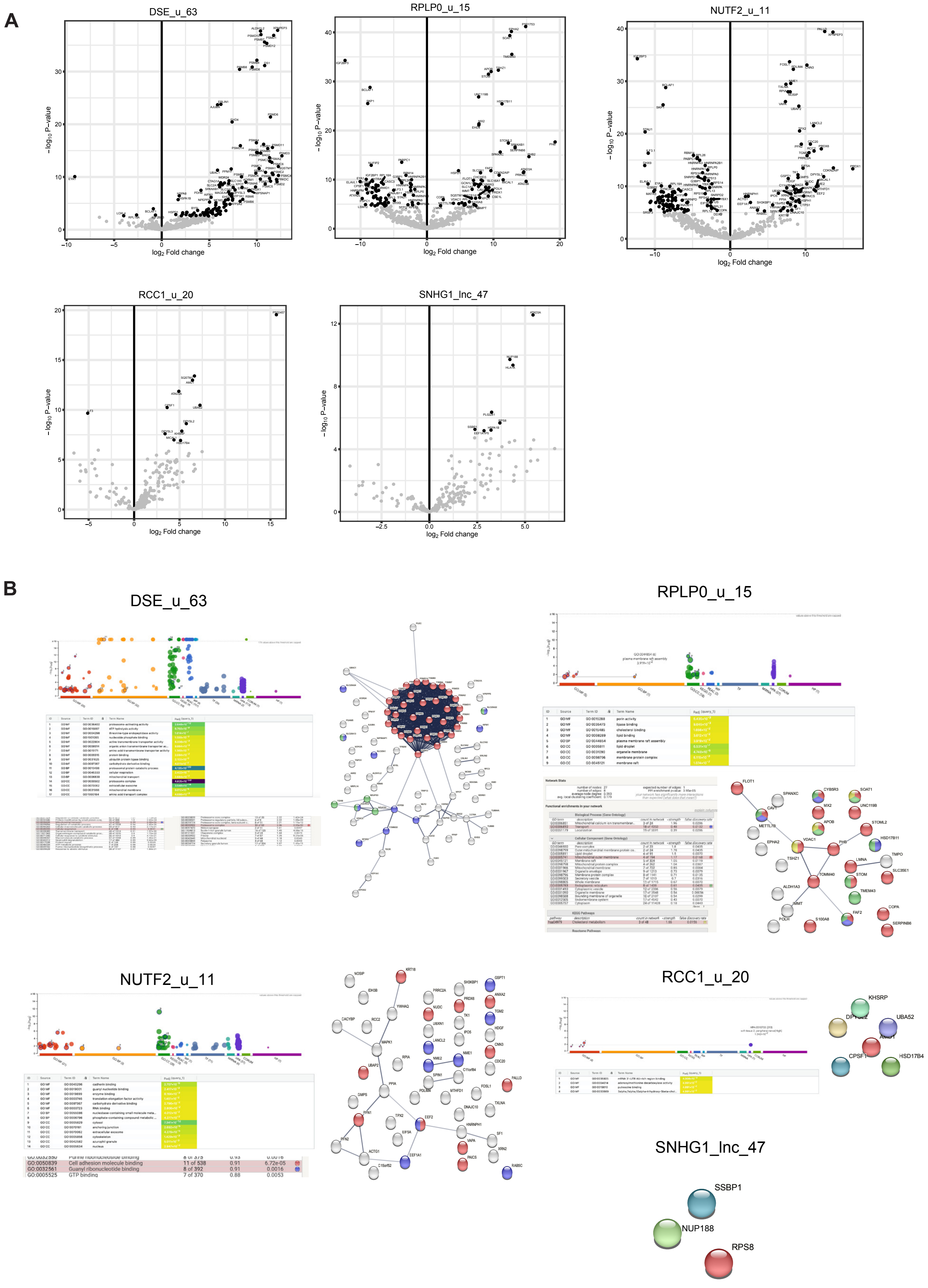


Supplementary figure 6: Further characterization of putative microprotein CO-IPs. A) Scatterplots of full DEP output for GFP-trap CO-IP with PBS washing conditions of 2 biological with 3 technical replicates for each condition. Prey had to be present in 6/6 replicates. Hits: LFC ≥ 1.5, DEP adjusted p-value < 0.05 (black), non-hits (grey). B) String database and gprofiler GO analysis output (where possible) for interaction partners identified in Fig.3 a-d.

### Supplementary figure 7 (related to Fig.3)

## A

##### DSE_u_63

MRVESSRGPLLMMAFTRTCRGFSPVSRCMTSKACLTMHTVMSFLPLLQPCIIMELVRRSTMGH

| Elm Name | Instances (Matched Sequence) | Positions | View in Jmol | Elm Description | Cell Compartment | Pattern | PHI-Blast Instance Mapping | Structural Filter Info | Probability |
|---|---|---|---|---|---|---|---|---|---|
| CLV_NRD_NRD_1 | RRS | 57-59 [A] | - | N-Arg dibasic convertase (NRD/Nardilysin) cleavage site (X-\|-R-K or R-\|-R-X). | extracellular, Golgi apparatus, cell surface | (.RK)\|(RR[^KR]) | - | - | 7.465e-03 |
| CLV_PCSK_KEX2_1 | RRS | 57-59 [A] | - | Yeast kexin 2 cleavage site (K-R-\|-X or R-R-\|-X). | extracellular, Golgi apparatus | [KR]R. | - | - | 7.973e-03 |
| CLV_PCSK_SKI1_1 | RCMTS | 27-31 [A] | - | Subtilisin/kexin isozyme-1 (SKI1) cleavage site ([RK]-X-[hydrophobic]-[LTKF]-\|-X). | endoplasmic reticulum lumen, endoplasmic reticulum, Golgi apparatus, extracellular | [RK].[AILMFV][LTKF]. | - | - | 6.821e-03 |
| CLV_Separin_Metazoa | ELVRR | 54-58 [A] | - | Separase cleavage site, best known in sister chromatid separation. | centrosome, nucleus, cytosol | E[IMPVL][MLVP]R. | - | - | 3.410e-04 |
| DEG_APCC_DBOX_1 | SRGPLLMMA | 6-14 [A] | - | An RxxL-based motif that binds to the Cdh1 and Cdc20 components of APC/C thereby targeting the protein for destruction in a cell cycle dependent manner | nucleus, cytosol | .R..L..[LIVM]. | - | - | 7.677e-04 |
| DEG_Nend_UBRbox_1 | MRVE | 1-4 [A] | - | N-terminal motif that initiates protein degradation by binding to the UBR-box of N-recognins. This N-degron variant comprises N-terminal Arg or Lys as destabilizing residue. | cytosol | ^M{0,1}[RK][^P]. | - | - | 2.064e-04 |
| DOC_CDC14_PxL_1 | MSFLPLLQP | 41-49 [A] | - | The PxL substrate docking motif enhances the Cdc14 phosphatase-substrate interaction and promotes subsequent dephosphorylation. | nucleus, nucleolus, cytosol | [FYLIM]..[YVILA]P.L.. | - | - | 2.446e-04 |
| DOC_WW_Pin1_4 | RGFSPV | 20-25 [A] | - | The Class IV WW domain interaction motif is recognised primarily by the Pin1 phosphorylation-dependent prolyl isomerase. | cytosol, nucleus | ...([ST])P. | - | - | 1.543e-02 |
| LIG_14-3-3_CanoR_1 | RCMTSKACL<br>RRSTMG<br>RSTMG | 27-35 [A]<br>57-62 [A]<br>58-62 [A] | -<br>-<br>- | Canonical Arg-containing phospho-motif mediating a strong interaction with 14-3-3 proteins. | nucleus, internal side of plasma membrane, cytosol | R[^DE]{0,2}[^DEPG]([ST])(([FWYLMV].)\|([^PRIKGN]P)\|([^PRIKGN].{2,4}[VILMFWYP])) | - | - | 4.477e-03 |
| LIG_LIR_Nem_3 | SFLPL | 42-46 [A] | - | Nematode-specific variant of the canonical LIR motif that binds to Atg8 protein family members to mediate processes involved in autophagy. | cytosol, cytoplasmic side of late endosome membrane | [EDST].{0,2}[WFY]..[ILVFY] | - | - | 6.362e-03 |
| LIG_MYND_1 | PLLQP | 45-49 [A] | - | PxLxP motif is recognized by a subset of MYND domain containing proteins. | nucleus | P.L.P | - | - | 6.499e-04 |
| LIG_PCNA_PIPBox_1 | MHTVMSFLPL | 37-46 [A] | - | The PCNA binding motifs include the PIP Box and the APIM motif, and are found in proteins involved in DNA replication, repair, methylation and cell cycle control. | nuclear replication fork, nucleus | [QM].[^FHWY][LIVM][^P][^PFWYMLIV](([FYHL][FYW])\|([FYH][FYWL]))... | - | - | 8.663e-05 |
| LIG_WRC_WIRS_1 | VMSFLP | 40-45 [A] | - | WRC interacting receptor sequence (WIRS) is a highly conserved and widespread interaction motif that is employed by diverse membrane proteins to recruit the WRC to initiate the dynamic rearrangements of the actin cytoskeleton. | cytosol, actin cytoskeleton, protein-containing complex | [FYILMV].[TS]F(G\|[^P]). | - | - | 5.802e-04 |
| MOD_CDK_SPxxK_3 | RGFSPVSR | 20-27 [A] | - | Longer version of the CDK phosphorylation site which shows specificity towards a lysine/arginine residue at position +4 after the phospho-Ser/Thr | cycloplasmic cyclin-dependent protein kinase holoenzyme complex, origin recognition complex, spindle, microtubule, cytosol, nucleus | ...([ST])P..[RK] | - | - | 1.929e-03 |
| MOD_CK1_1 | SPVSRCM | 23-29 [A] | - | CK1 phosphorylation site | cytosol, nucleus | S..([ST])... | - | - | 1.704e-02 |
| MOD_GSK3_1 | SPVSRCMT | 23-30 [A] | - | GSK3 phosphorylation recognition site | cytosol, nucleus | ...([ST])...[ST] | - | - | 2.679e-02 |
| MOD_NEK2_2 | FTRTCRG | 15-21 [A] | - | The more tolerant version of the Nek2 phosphosite motif | centrosome, Ndc80 complex, condensed nuclear chromosome outer kinetochore, cytosol, nucleus, kinetochore, centriole, microtubule organizing center | [FLMW][^P][^P]([ST])[^PDECAN][RKH]. | - | - | 1.389e-03 |
| MOD_PKA_1 | RRSTMGH | 57-63 [A] | - | Main preference for PKA-type AGC kinase phosphorylation. | cAMP-dependent protein kinase complex, cytosol, nucleus | [RK][RK].([ST])[^P].. | - | - | 2.315e-03 |
| MOD_PKA_2 | VRRSTMG<br>RRSTMGH | 56-62 [A]<br>57-63 [A] | -<br>- | Secondary preference for PKA-type AGC kinase phosphorylation. | cytosol, nucleus, cAMP-dependent protein kinase complex | .R.([ST])[^P].. | - | - | 9.458e-03 |
| MOD_Plk_4 | TMHTVMS<br>TVMSFLP | 36-42 [A]<br>39-45 [A] | -<br>- | Ser/Thr residue phosphorylated by Plk4 | nucleus, cytosol, SCF ubiquitin ligase complex, cleavage furrow, centriole, gamma-tubulin ring complex, centriolar satellite, pericentriolar material | ..[^IRFW]([ST])[ILMVFWY][ILMVFWY]. | - | - | 6.019e-03 |
| MOD_ProDKin_1 | RGFSPVS | 20-26 [A] | - | Proline-Directed Kinase (e.g. MAPK) phosphorylation site in higher eukaryotes. | cytosol, nucleus | ...([ST])P.. | - | - | 1.543e-02 |
| TRG_DiLeu_BaLyEn_6 | SRGPLL<br>SFLPLL | 6-11 [A]<br>42-47 [A] | -<br>- | Dileucine motifs lacking Glu+1 with Pro-Arg preference at +4 sorting in Endosomal-Basolateral-Lysosomal trafficking. | plasma membrane, cytosol, cytoplasmic side of membrane | [^E]..[RP]L[LI] | - | - | 1.053e-03 |
| TRG_ER_diArg_1 | VRR | 56-58 [A] | - | The di-Arg ER retention motif is defined by two consecutive arginine residues (RR) or with a single residue insertion (RXR). The motif is completed by an adjacent hydrophobic/arginine residue which may be on either side of the Arg pair. | cytosol, endoplasmic reticulum membrane, integral protein, ER-Golgi transport vesicle membrane, endoplasmic reticulum membrane, Golgi-ER transport vesicle membrane, rough endoplasmic reticulum, endoplasmic reticulum cisterna | ([LIVMFYWPR]R[^YFWDE]{0,1}R)\|(R[^YFWDE]{0,1}R[LIVMFYWPR]) | - | - | 5.369e-03 |

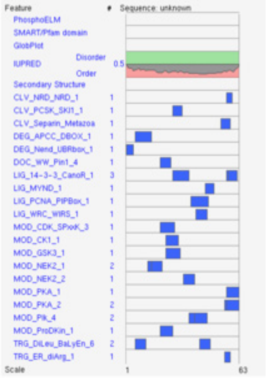


##### RCC1_u_20

MCKILEEDSRERERDQRSQG

| Elm Name | Instances (Matched Sequence) | Positions | View in Jmol | Elm Description | Cell Compartment | Pattern | PHI-Blast Instance Mapping | Structural Filter Info | Probability |
|---|---|---|---|---|---|---|---|---|---|
| DEG_Nend_UBRbox_4 | MCK | 1-3 [A] | - | N-terminal motif that initiates protein degradation by binding to the UBR-box of N-recognins. This N-degron variant comprises N-terminal Cys as destabilizing residue. | cytosol | ^M{0,1}(C). | - | - | 1.768e-05 |
| DOC_PP2A_B56_1 | MCKILE | 1-6 [A] | - | Docking site required for the regulatory subunit B56 of PP2A for protein dephosphorylation. | nucleus, kinetochore, chromosome, centromeric region, cytosol | ([LMFYWIC]..I.E)\|(L..[IVLWC].E). | - | - | 1.458e-03 |

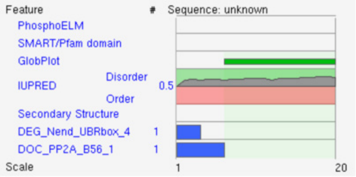


##### RPLP0_u_15

MGVGVTRPELPVSPQ

| Elm Name | Instances (Matched Sequence) | Positions | View in Jmol | Elm Description | Cell Compartment | Pattern | PHI-Blast Instance Mapping | Structural Filter Info | Probability |
|---|---|---|---|---|---|---|---|---|---|
| DOC_WW_Pin1_4 | LPVSPQ | 10-15 [A] | - | The Class IV WW domain interaction motif is recognised primarily by the Pin1 phosphorylation-dependent prolyl isomerase. | cytosol, nucleus | ...([ST])P. | - | - | 1.543e-02 |
| LIG_FHA_2 | GVTRPEL | 4-10 [A] | - | Phosphothreonine motif binding a subset of FHA domains that have a preference for an acidic amino acid at the pT+3 position. | nucleus, Replication fork | ..(T)..[DE]. | - | - | 8.286e-03 |
| LIG_SH3_3 | GVGVTRP<br>VTRPELP<br>PELPVSP | 2-8 [A]<br>5-11 [A]<br>8-14 [A] | -<br>-<br>- | This is the motif recognized by those SH3 domains with a non-canonical class I recognition specificity | plasma membrane, focal adhesion, cytosol | ...[PV]..P | - | - | 1.317e-02 |
| MOD_CK2_1 | VGVTRPE | 3-9 [A] | - | Casein kinase 2 (CK2) phosphorylation site | protein kinase CK2 complex, nucleus, cytosol | ...([ST])..E | - | - | 1.457e-02 |
| MOD_NMyristoyl | MGVGVTR | 1-7 [A] | - | Generic motif for N-Myristoylation site. | anchored to plasma membrane, cytosol | ^M{0,1}(G)[^EDRKHPFYW]..[STAGCN][^P] | - | - | 2.502e-05 |

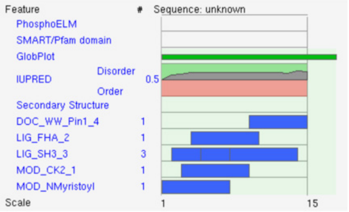


##### SNHG1_lnc_47

MFSPQELTGEGMGQDPSLCKASVTVMFQVFLMTVCLYLTEQLVAIYH

| Elm Name | Instances (Matched Sequence) | Positions | View in Jmol | Elm Description | Cell Compartment | Pattern | PHI-Blast Instance Mapping | Structural Filter Info | Probability |
|---|---|---|---|---|---|---|---|---|---|
| DEG_Nend_Nbox_1 | MFS | 1-3 [A] | - | N-terminal motif that initiates protein degradation by binding to the N-box of N-recognins. This N-degron variant comprises N-terminal a bulky hydrophobic residue as destabilizing residue. | cytosol | ^M{0,1}[FYLIW][^P] | - | - | 2.302e-04 |
| LIG_TRAF2_1 | SPQE | 3-6 [A] | - | Major TRAF2-binding consensus motif. Members of the tumor necrosis factor receptor (TNFR) superfamily initiate intracellular signaling by recruiting the C-domain of the TNFR-associated factors (TRAFs) through their cytoplasmic tails. | cytosol | [PSAT].[QE]E | - | - | 4.300e-03 |

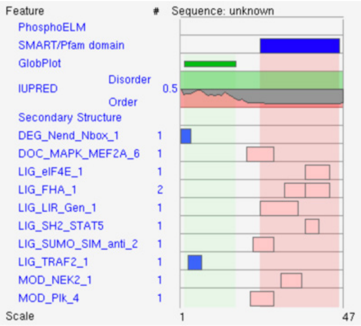


## B

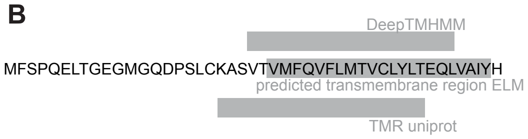


Supplementary figure 7: Motif search for each of the putative microproteins. A) ELM motif search output for each of the putative microproteins (where possible). B) Predicted transmembrane region for SNHG1_lnc_47 by different algorithms/databases.

### Supplementary Figure 8 (related to Fig.4)

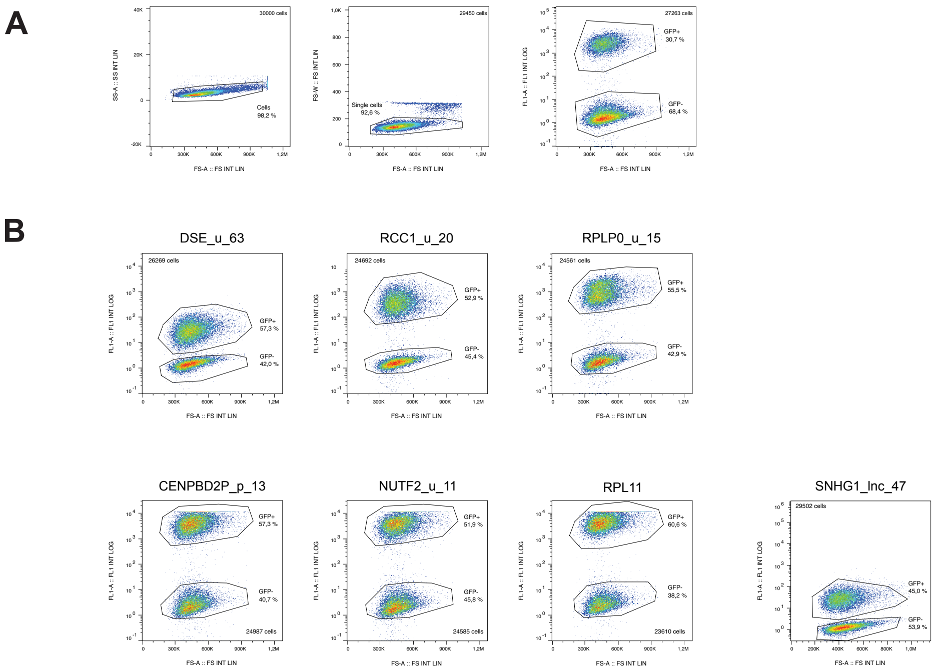


Supplementary Figure 8: Examples of flow cytometry gatings for the microprotein rescue trials. A) Illustrative example of gating workflow to extract GFP- positive and GFP-negative single cells populations. Shown is a coculture of A375 Cas9 GFP cells and DSE_u_63 untagged rescue cells lines one day after transfection with TRAC sgRNA. B) Examples of gating for GFP-positive and GFP-negative single cell populations for co-cultures of A375 Cas9 cells with the respective GFP-tagged microprotein rescue cells one day after transfection with TRAC sgRNA.

### Supplementary Figure 9 (related to Fig.5)

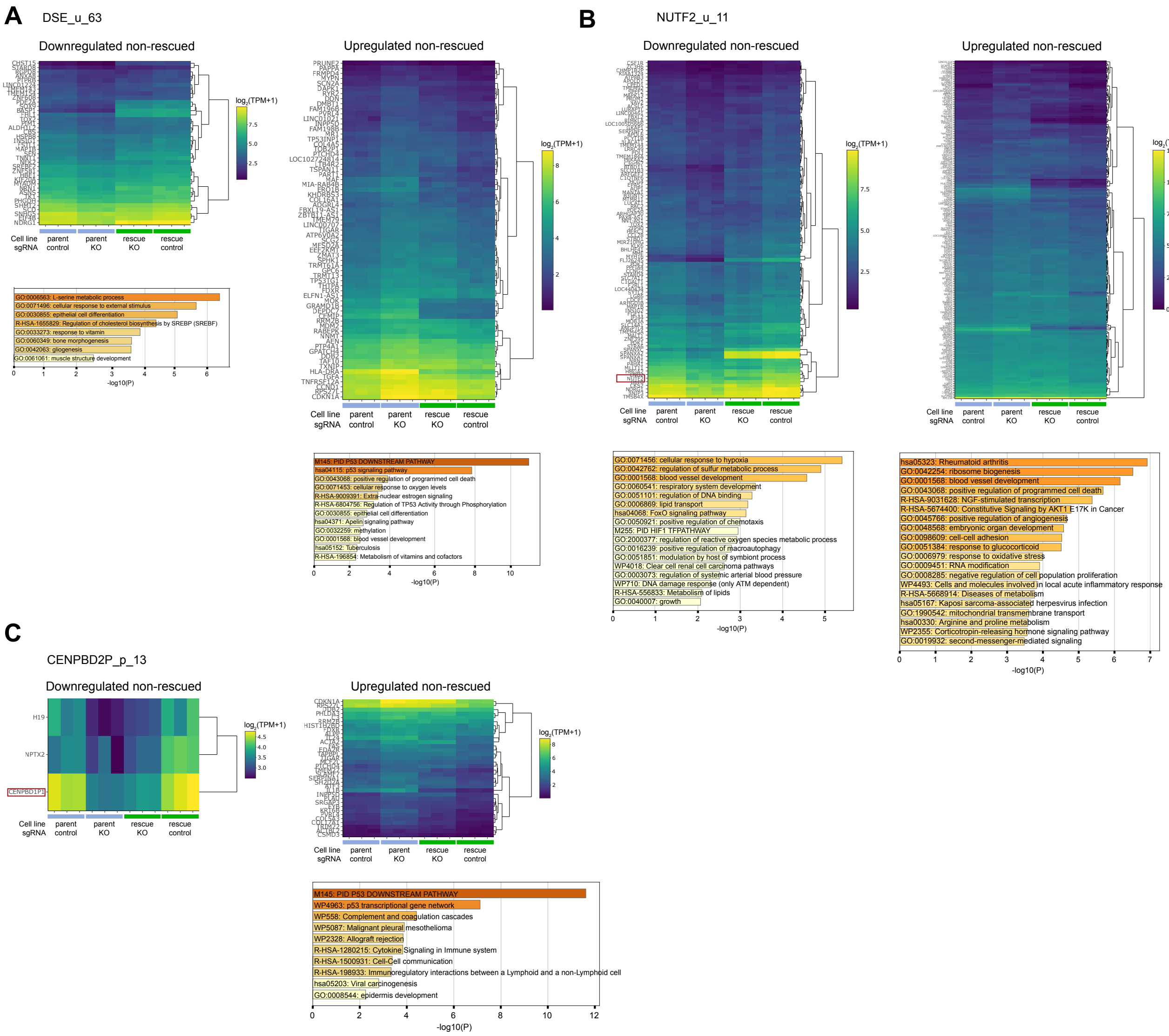


Supplementary figure 9: RNA-seq rescue experiments with GFP-tagged codon-altered putative microproteins. A-C) left panel: Heatmap of genes that were downregulated upon treatment of A375 Cas9 cells with the respective ORF sgRNA and that could not be rescued in microprotein-GFP expressing A375 Cas9 cells. The respective sORF gene is highlighted in red. Bargraph of Metascape GO term analysis (where possible). right panel: Heatmap of genes that were upregulated upon treatment of A375 Cas9 cells with the respective ORF sgRNA and that could not be rescued in microprotein-GFP expressing A375 Cas9 cells. Bargraph of Metascape GO term analysis. For all panels: Hits: genes displaying an average (of 2 replicates) LFC of ≥ 0.5 or ≤ –0.5 between microprotein sgRNA and TRAC sgRNA treated A375 Cas9 cells and showing an average expression variability of ≤10% between the TRAC conditions (microprotein-GFP vs. parental cells) and an DeSeq2 adjusted p-value of ≤ 0.05 , Rescued hits: same as hits but additionally showing a difference in LFC of ≥ 0.5 between microprotein-GFP and parental cell lines.

### Supplementary Figure 10  (related to Fig.5)

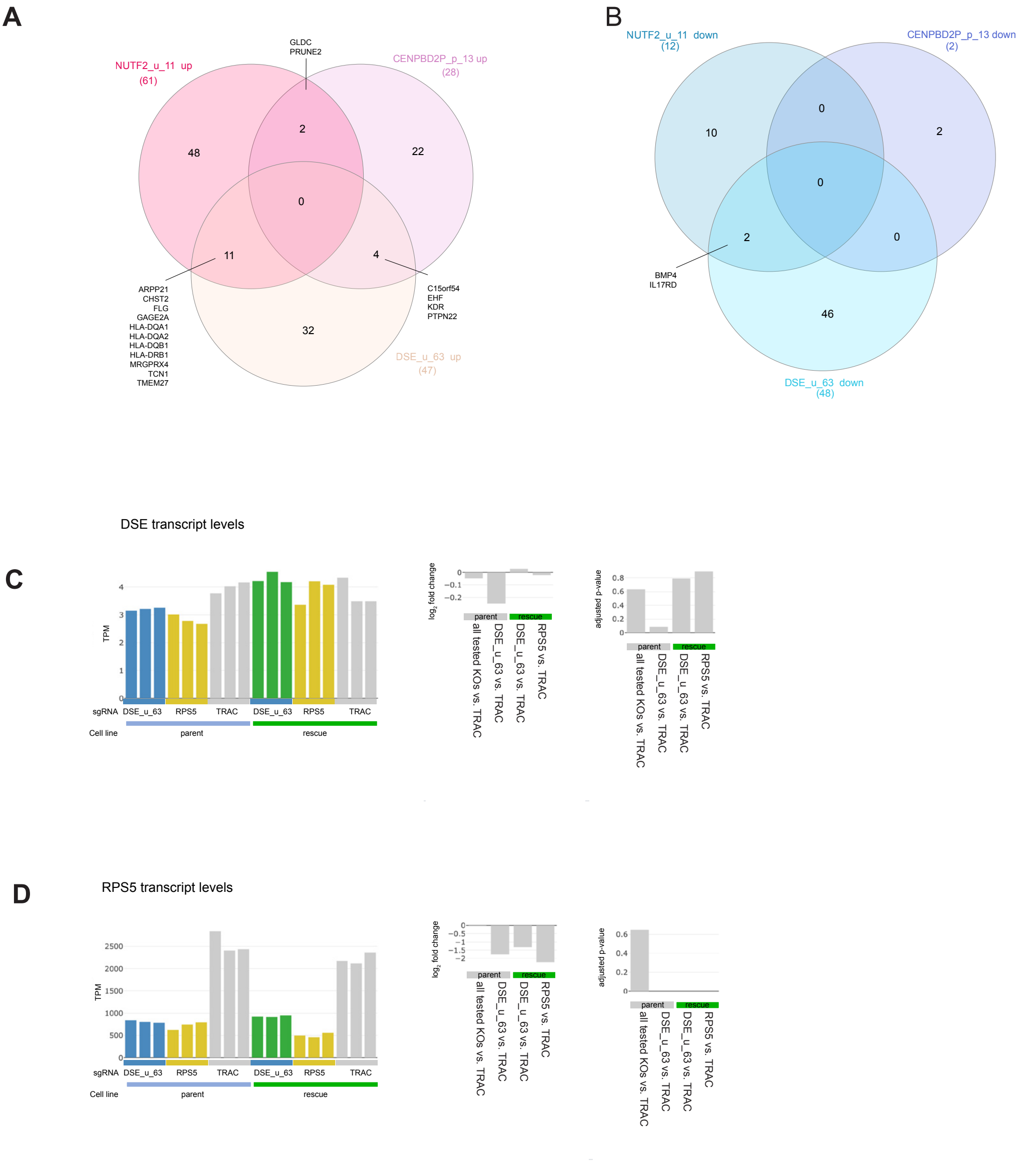


Supplementary figure 10: Overlap between rescued genes of different microprotein knockouts. a-b) Venn-diagrams depicting overlap between upregulated rescued genes (A) and downregulated rescued genes (B) of the different microprotein sgRNA conditions. C) Bargraph of DSE transcript levels across the different tested conditions from RNA-seq output and associates bargraphs indicating LFC and adjusted p-value. D) Same as C but for RPS5 transcript levels.
